# Estimating the redox state of the plastoquinone pool in algae and cyanobacteria *via* OJIP fluorescence: perspectives and limitations

**DOI:** 10.1101/2025.08.05.668659

**Authors:** Tomáš Zavřel, Anne-Christin Pohland, Tobias Pfennig, Anna Barbara Matuszyńska, Szilvia Z. Tóth, Gábor Bernát, Jan Červený

## Abstract

The redox state of the plastoquinone pool (PQ-redox) acts as a central element in a variety of intracellular signal pathways. Several methods for determining PQ-redox have been established. Although some of these methods may be quantitative, such as those based on liquid chromatography, they are typically sensitive to sample preparation. Here, we critically evaluate the use of fast chlorophyll *a* fluorescence induction kinetics (the so-called OJIP transient) for semi-quantitative PQ-redox estimation in green algae (*Chlorella vulgaris*) and cyanobacteria (*Synechocystis* sp. PCC 6803). The method, based on the evaluation of relative fluorescence yield at the J-step of the OJIP transient (VJ, VJ’), has already been reported; however, thus far, it has been used mostly for studying dark-acclimated leaves, which limits its range of application. Here, we show that the OJIP transient can be used for semi-quantitative estimation of PQ-redox in algal and cyanobacterial cell cultures, in addition to plants. We further show that it can reflect PQ-redox in both dark-acclimated and light-acclimated samples. Our systematic comparison of Multi-Color PAM, AquaPen, and FL 6000 fluorometers demonstrates that accurate measurement of VJ and VJ’ parameters in suspension cultures requires low culture density and a high-intensity saturation pulse. We further show that with increasing light intensity to which the cells are exposed, the state of photosystem II (PSII) changes due to light-induced reduction of quinone A (QA-) and conformational changes, which in turn influence both the sensitivity and dynamic range of the VJ’ parameter towards PQ-redox estimation. A comparison of fluorescence transients in *Chlorella* and *Synechocystis* revealed high homeostatic control over PQ-redox in *Synechocystis*, maintained by terminal oxidases present at the thylakoid membrane. While we discuss certain limitations, our systematic assessment suggests that the OJIP method has great potential to become a routine tool for semi-quantitative PQ-redox estimation under a wide range of experimental conditions in green algae and cyanobacteria.

## Background

Photosynthesis is a fundamental process on Earth that provides energy (and produces biomass) for almost the entire food chain. During photosynthesis, light energy is captured by light-harvesting antennae. The captured energy is ultimately transferred to the reaction centers (special pair of chlorophyll molecules) of the photosystems where charge separation takes place (Nelson and Yocum 2006). Using the positive charges created, water is split into molecular oxygen and protons at the oxygen evolving complex of photosystem II (PSII) (Li et al. 2024), while the released electrons are passed through the photosynthetic electron transport chain (PETC) towards photosystem I (PSI) and to the terminal electron acceptors such as NADP^+^. The mobile, membrane-bound photosynthetic electron transport chain (PETC) electron carrier, plastoquinone (PQ), accepts electrons from PSII, and transfers them (in the form of plastoquinol, PQH2) primarily to cytochrome *b*6/*f* (Cyt *b*6/*f*), from which they are further transferred to PSI through water soluble electron carriers, plastocyanin or cytochrome *c*6 (Blankenship 2002). The redox state of the PQ pool (PQ-redox hereafter), i.e. the portion of PQ oxidized, relative to the portion of PQ reduced, has an important signaling function, triggering a variety of metabolic processes, including light acclimation, biosynthesis of metabolites, or gene expression (Havaux 2020).

The PQ pool can be reduced and oxidized by several processes. Besides linear electron flow, these include cyclic electron flow around PSI (PSI-CEF), chlororespiration, and water-water cycle (Miyake 2010; Bernát and Rögner 2011). In addition, the PQ pool can be oxidized by terminal oxidases (TOs), including plastid terminal oxidase (PTOX) (Joët et al. 2002). In cyanobacteria, the PQ pool can be further reduced by respiratory components present at the thylakoid membrane (TM) (Lea-Smith et al. 2013). PQ-redox is an important element of stress acclimation, regulating state transitions (Calzadilla and Kirilovsky 2020; Virtanen and Tyystjärvi 2023), synthesis of carotenoids, expression of PSII and PSI genes, proteins from chloroplast and nucleus, superoxide dismutase, and many other enzymes (Mattila et al. 2020; Havaux 2020; Pilarska et al. 2023). Furthermore, PQ-redox plays a key role in mitigating heat and salt stress (Pshybytko et al. 2008; Pilarska et al. 2023), acclimation to high light exposure (Lepetit et al. 2013) or to changing light quality (Bernát et al. 2021; Zavřel et al. 2024), and in the establishment of plant response mechanisms to pathogen infection (Nosek et al. 2015). PQ can also serve as an antioxidant (Khorobrykh and Tyystjärvi 2018).

PQ and PQH2 can be quantified by HPLC (Kruk and Karpinski 2006). If the extraction is performed quickly enough to preserve the native state, this method offers fully quantitative results, and can further discriminate photochemically active PQ within the total PQ pool. With a proper arrangement, it can be used to determine PQ-redox dependence on various conditions, as well as to follow PQ-redox shifts, e.g., upon changes in light quality or light intensity (Mattila et al. 2020). However, despite all the advantages, this method faces limitations in detecting fast or transient PQ-redox changes. For this reason, optical methods have been developed, based either on light absorbance or chlorophyll *a* (Chl *a*) fluorescence (Berry et al. 2002; Tóth et al. 2007; Tsimilli-Michael et al. 2009; Fukunaga et al. 2024; Naydov et al. 2024). The absorbance measured at 254 nm or 263 nm can detect fast PQ-redox changes upon dark acclimation following illumination (Schmidt-Mende and Witt 1968; Ivanov et al. 2007). However, the use of UV-Vis spectrophotometry for *in vivo* applications remains limited due to generally demanding setup. All Chl *a* fluorometers, on the other hand, typically include a source of actinic light (AL), making them more applicable in studying light-dependent PQ-redox shifts. Simultaneous measurement of Chl *a* fluorescence and 820 nm transmission has also been proven effective for detecting light-dependent PQ-redox dynamics (Schansker et al. 2005).

Indeed, the use of Chl *a* fluorescence for PQ-redox monitoring possesses certain limitations. Here, we provide a systematic evaluation of the use of fast Chl *a* fluorescence induction, the so-called OJIP transient (Strasser and Govindjee 1992; Stirbet et al. 2018), towards PQ-redox determination under a variety of conditions. The fluorescence transient originates from radiative deactivation of gradually excited PSII during illumination by a saturation pulse (SP). De-excitation of PSII involves multiple intermediate steps of reduction and oxidation of quinone A (QA) and quinone B (QB), each characterized by rate constants generally spanning over several orders of magnitude, approximately between 10^1^ - 10^9^ s^−1^ (Lazár 2003). Since the SP length typically ranges between 600 - 2 000 ms, the redox equilibrium within PSII during SP is further influenced by other PETC components. The various phases of the OJIP transient thus reflect kinetic bottlenecks of the electron transport chain (Schansker et al. 2005), including the exchange of a reduced PQ molecule to an oxidized one at the QB site (J level, at around 2 ms), without involvement of downstream PETC components. The I level (at around 30 ms) represents the limitation in the reoxidation of PQH2 by the Cyt *b*6/*f* complex. The P level, appearing typically at 200-300 ms in dark-acclimated cells, represents the transient block imposed by inactive FNR. The whole OJIP curve thus reflects activity of the entire PETC up to the Calvin-Benson-Bassham cycle (CBB) (Schansker et al. 2005; Tsimilli-Michael 2020).

The fluorescence yield at the first inflection point J (FJ) has been identified as the most promising parameter for the PQ-redox estimation (Tóth et al. 2007). Later on, the normalized FJ value, VJ (Eq. 1), has been recommended as a more precise PQ-redox proxy, since it considers both the initial (FO) and maximal (FM) fluorescence yields (Tsimilli-Michael et al. 2009). In this work, we first examined the applicability of the VJ parameter for assessing PQ-redox state in representative strains of green algae (*Chlorella vulgaris* ATCC 30821, *Chlorella* hereafter) and cyanobacteria (*Synechocystis* sp. PCC 6803, *Synechocystis* hereafter) in a dark-acclimated state. Building on these results, we extended the approach to estimate PQ-redox under light-acclimated conditions, by using parameter VJ’ (Eq. 2) (Tsimilli-Michael et al. 2009). A systematic comparison of three fluorometers (Multi-Color PAM, AquaPen, and FL 6000) showed that both VJ and VJ’ parameters can estimate PQ-redox well in both species when certain assumptions are met. These include fully or at least mostly opened PSII centers (in order to keep the fraction of reduced Q_A_ ^−^ low), strong SP in order to reduce all Q_A_, low culture density to avoid fluorescence scattering and reabsorption, and correct identification of FJ timing. When these criteria were satisfied, both VJ and VJ’ reflected PQ reduction reliably during various perturbations: under actinic light (AL) compared with darkness and under inhibition of CBB cycle and TOs with glycolaldehyde and potassium cyanide, respectively - consistently with previous reports (Khorobrykh et al. 2020; Fukunaga et al. 2024). VJ also reflected PQH2 oxidation after the addition of methylviologen (MV) (Schansker et al. 2005). The method holds strong potential to become a widely used tool for non-invasive, semi-quantitative PQ-redox determination in algae and cyanobacteria cultures - expanding its application beyond plants, where it has already been extensively used.

## Methods

### Fluorometer setup and VJ calculation

Fast fluorescence induction kinetics (OJIP transients) were measured using three fluorometers: Multi-Color PAM (Walz, Germany), AquaPen and FL 6000 (Photon System Instruments, Czechia). During the tests examining the interplay between culture density and SP intensity, the photon flux density (PFD) of the blue SP (440-460 nm) varied between 1 100 and 7 500 µmol photons m^−2^ s^−1^ for *Chlorella*, and the PFD of the red SP (623-630 nm) ranged from 975 to 6 500 µmol photons m^−2^ s^−1^ for *Synechocystis*. In all other experiments, OJIP transients were measured by Multi-Color PAM and both SP wavelength and intensity were kept constant: 440 nm and 2 244 µmol photons m^−2^ s^−1^ for *Chlorella* and 625 nm and 1 904 µmol photons m^−2^ s^−1^ for *Synechocystis*.

AquaPen and FL 6000 do not employ a measuring light (ML). In contrast, in Multi-Color PAM low-intensity ML (PFD < 2 µmol photons m^−2^ s^−1^) was used during SP application, during acclimation of *Chlorella* (440 nm ML) and *Synechocystis* (625 nm ML) cultures under AL, as well as ∼1 s before SP application in dark-acclimated cultures not treated with DCMU. In the Multi-Color PAM, stirring was applied during the whole incubation period, and the stirring was interrupted ∼1 s before each OJIP measurement. The OJIP fluorescence recording was initiated 10 ms prior to the SP application. The SP was applied as a multiple turnover flash, with the ML modulation frequency set to 100 kHz. At the onset of the SP, the sample-and-hold feature of the Multi-Color PAM, keeping the original analog signal for a short time to allow digitization, was disabled. Detailed settings of all fluorometers used in the experiments are summarized in Supplementary Table S1. The emission spectra of the LEDs employed in each fluorometer are shown in Supplementary Figure S1.

To follow the dynamics of the redox state of the PQ pool in fully dark-acclimated cultures, the parameter VJ was derived from OJIP curves double-normalized between FO and FM (Tóth et al. 2007):

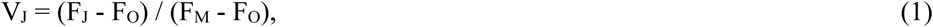

where FJ is the fluorescence intensity at the J point of the OJIP transient (see below for details), FO is the initial fluorescence yield and FM is the maximal fluorescence yield. To determine FJ, the fluorescence signal between 0.5 and 5 ms was fitted with a ninth-degree polynomial function (Akinyemi et al. 2023). The algebraic derivatives of this fitted function were then used to identify the inflection points as the roots of the second derivative with positive values in the third derivative. The inflection point closest to the expected 2 ms timing was assigned FJ.

To probe PQ-redox in light-acclimated or partially dark-acclimated cultures, the parameter VJ’ was used (Tsimilli-Michael et al. 2009):

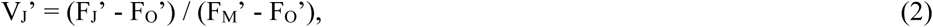

where FO’, FJ’ and FM’ denote the initial fluorescence, the fluorescence at the J point, and the maximal fluorescence, respectively, measured in light-acclimated or partially dark-acclimated samples. The timing of FJ’ was again identified as the inflection point of the reconstructed fluorescence nearest to 2 ms.

### Strains and cultivation conditions

Stock cultures of *Synechocystis* sp. PCC 6803 and *Chlorella vulgaris* ATCC 30821 were cultivated in Erlenmeyer flasks in BG-11 medium (Rippka et al. 1979; van Alphen et al. 2018) at 25°C, under 25 µmol photons m^−2^ s^−1^ of white light, in ambient air. For the comparative experiments, both strains were cultivated in Multi-Cultivator MC-1000-MIX (Photon System Instruments, Czechia) in a turbidostat regime. The culture density was maintained between OD680 = 0.395 - 0.405, corresponding to ∼0.54 and ∼0.69 mg Chl *a* L^−1^ for *Synechocystis* and *Chlorella*, respectively. In Multi-Cultivators, the cultures were kept under 25 µmol photons m^−2^ s^−1^ of warm-white LEDs (Supplementary Figure S1), at 25°C, and aerated either with ambient air or with air enriched to 0.5% (v/v) CO2 using Gas Mixing System GMS-150, Photon System Instruments. The aeration rate was maintained at approximately 30 mL min^−1^.

### Experimental setup

#### Fluorometer comparison under varying culture density and SP intensity

Cultures were withdrawn from Erlenmeyer flasks, centrifuged (3 000 x *g*, 10 min), and resuspended in fresh BG-11 medium to prepare a series of cultures with concentrations of 0.1 - 24.6 mg Chl *a* L^−1^. The interplay between culture density and SP intensity was measured using Multi-Color PAM, AquaPen and FL 6000 fluorometers, and dark-acclimated cultures. To avoid detector saturation, OJIP curves were first recorded with the maximal SP intensity in both Multi-Color PAM and FL 6000 fluorometers throughout all culture densities. After the culture density providing the highest signal was identified (3.8 and 2.7 mg Chl *a* L^−1^ for *Synechocystis* and *Chlorella*, respectively; Supplementary Figure S2), detector gain in Multi-Color PAM and FL 6000 was set such that the fluorescence signal did not exceed 80% of the detector range under maximal SP intensity.

In the most recent AquaPen version, detector settings cannot be adjusted. All cultures were therefore measured with default settings, leading to detector saturation in many measurements (Fig. 1, Supplementary Figures S2-S3). Details of the settings of the fluorometers are provided in Supplementary Table S1.

**Fig. 1.**
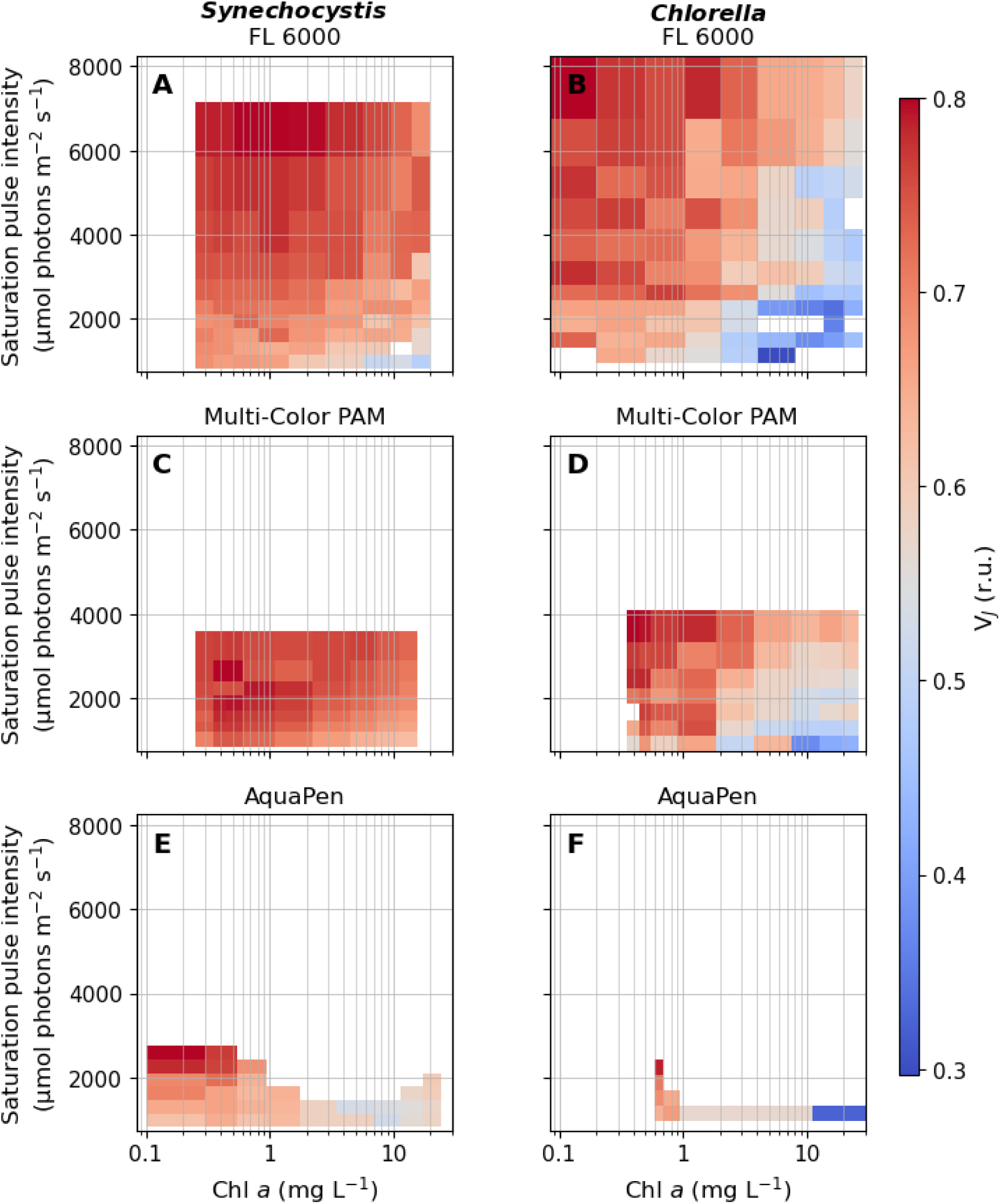
Heat maps of the VJ parameter derived from OJIP transients (Supplementary Figure S2-S3) recorded by FL 6000 (A-B), Multi-Color PAM (C-D), and AquaPen (E-F) fluorometers across a matrix of SP intensities (975-7 500 µmol photons m^−2^ s^−1^) and culture densities (0.1 to 24.6 mg Chl *a* L^−1^). Prior to the measurements, *Synechocystis* (A, C, E) and *Chlorella* (B, D, F) cultures were concentrated by centrifugation, suspended in fresh BG-11 medium to final cell densities and dark-acclimated (see Materials and Methods for details). The timing of FJ was determined by identifying inflection points within the 0.6-25 ms range, as marked by the grey rectangles in Supplementary Figure S3. For better visualization, the x-axis is shown on a logarithmic scale. Missing values for FL 6000 fluorometer resulted either from high noise level under the lowest SP intensities compromising reliable FJ identification, or from identifying the first inflection point outside of the specified range. For AquaPen, missing data resulted from detector saturation; when the saturation occurred, the OJIP curves were excluded from analysis. A complete description of the settings of all used fluorometers, including SP intensities used, is provided in Supplementary Table S1.

#### Treatment with DCMU (3-(3,40-dichlorophenyl)-1,1-dimethylurea)

The effect of DCMU was tested on cultures dark-acclimated for 20 min. After withdrawing from the Multi-Cultivator, aliquots of DCMU stock solution (10 mM, dissolved in DMSO) were added to the 1.5 mL culture samples to reach final DCMU concentrations 0.1 nM - 20 μM. As controls, cultures without DCMU addition were used. To avoid QA- accumulation in PSII prior to the OJIP measurement, the measuring light in Multi-Color PAM was turned off before the OJIP measurements, and it was applied only during SP application.

#### Dark-to-light transition

1.5 mL culture aliquots were withdrawn from the Multi-Cultivator, dark-acclimated for 20 min and transferred to the cuvette of Multi-Color PAM under dim ambient light. Fast fluorescence induction kinetics was measured ∼30 s after the transition to the Multi-Color PAM cuvette holder. After the first OJIP measurement, white AL (Supplementary Figure S1) of PFD 100 µmol photons m^−2^ s^−1^ was turned on, and the OJIP transients were further measured in 30 s intervals for 5 min. Approximately 1 s before each SP, AL was turned off, and it was resumed immediately after the OJIP measurement. ML was kept on during the whole measurement course, starting ∼5 s before the first OJIP transient recorded in darkness. Multi-Color PAM settings are summarized in Supplementary Table S1. In experiments with methyl viologen (98%, Sigma-Aldrich, USA), the cultures were incubated in the presence of MV in dark at 25°C for 20 min before the measurements. MV was used in the final concentration 1 mM and 0.25 mM; the 1 M stock solution was obtained by dissolving MV in double deionized water.

#### Effect of GA and KCN under illumination

In experiments where the effect of glycolaldehyde (GA; Carl Roth, Germany) and potassium cyanide (KCN; ≥98%, Sigma-Aldrich, USA) was tested, each inhibitor was added to light-acclimated cultures. After withdrawing from the Multi-Cultivator, 1.5 mL culture aliquots were transferred to the cuvette of Multi-Color PAM, to acclimate under white AL (Supplementary Figure S1) of PFD 100 µmol photons m^−2^ s^−1^ for 10 min. OJIP transients were first measured without the presence of inhibitors. Right after this first OJIP measurement, either GA or KCN were added at final concentrations 25 mM and 1 mM, respectively, and OJIP transients were further measured in 30 s intervals for 5 min under illumination.

#### High light treatment

During transition from low to high light, 1.5 mL culture aliquots were withdrawn from the Multi-Cultivator, transferred to the cuvette of Multi-Color PAM and acclimated under white AL (Supplementary Figure S1) of PFD 25 µmol photons m^−2^ s^−1^ for 5 min. OJIP transients were first measured under 25 µmol photons m^−2^ s^−1^. Right after the first measurement, PFD was increased to 1 500 µmol photons m^−2^ s^−1^ and OJIP curves were further measured in 30 s intervals for 5 min under this high light.

## Results

### Technical aspects of the OJIP measurements and accurate determination of VJ

#### Fluorometer selection

In this work, the applicability of fast Chl *a* fluorescence rise (OJIP transient) was evaluated for estimating the redox state of the PQ pool in *Synechocystis* and *Chlorella*. As a PQ-redox proxy, the normalized value of the fluorescence signal at the J point (VJ and VJ’, Eq. 1-2) was selected. The fluorescence induction curves were recorded using a Multi-Color PAM fluorometer (Heinz Walz GmbH, Germany). This instrument uses modulated light (measuring light, ML) of low intensity to induce and monitor fluorescence changes. It can separate fluorescence signal induced by ML from that induced by AL or SP (Schreiber 1986). Typically, ML excites only a small fraction of the PSII reaction centers. This is important to understand, since VJ and VJ’, according to earlier results, reflect redox equilibrium within PSII - possibly between the reduced quinone A (QA-) and quinone B (QB) (Tomek et al. 2001; Schansker and Strasser 2005). Under normal conditions, when QA- re-oxidation *via* forward electron transport is not impaired, the effect of ML on the QA-/QB equilibrium is negligible. However, ML can affect the PSII redox balance significantly and even fully close PSII under specific conditions, such as in the presence of DCMU (Tóth et al. 2005).

The OJIP transients were also recorded by AquaPen and FL 6000 fluorometers. The principle of fluorescence measurement in these so-called PEA-type fluorometers not employing ML (named after Handy PEA fluorometer, Hansatech Instruments Ltd., UK (Kalaji et al. 2014)) is different from Multi-Color PAM. In PEA-type fluorometers, SP is the only light source, the excitation light and Chl *a* fluorescence are separated by filters, and the measured fluorescence is directly proportional to the SP intensity applied. In these fluorometers, comparing fluorescence traces across SP intensities requires initial normalization according to the SP intensity (Schansker et al. 2006). Differences between the tested fluorometers further include data sampling strategies and detector settings options. Data sampling in Multi-Color PAM is linear along the entire OJIP transient, whereas in both AquaPen and FL 6000 the fluorescence recording is non-linear within specific, predefined intervals. During our experiment comparing different fluorometers (Fig. 1, Supplementary Figures S2-S3), the initial 630 ms of the OJIP transients were recorded with varying temporal resolution: AquaPen, FL 6000, and Multi-Color PAM captured 63, 827 and 63 997 data points, respectively. Detector settings, including gain and offset, are accessible only in FL 6000 and Multi-Color PAM. In FL 6000, offset shifts the signal to lower or even negative voltage values, whereas in Multi-Color PAM it subtracts background signals. In addition, Multi-Color PAM offers the option to modify the detector time constant through Frequency and Damping settings. Lowering the detector time constant can become useful during slow kinetic measurement, however, for OJIP experiments, it is necessary to keep Damping low to secure the fastest measuring frequency and highest time resolution possible.

All tested fluorometers provided both blue and red SPs (Supplementary Figure S1) and detected the fluorescence signal in the red/far red part of the light spectrum (665-750 nm, Supplementary Table S1). The absolute values of the VJ parameter varied slightly among the three tested fluorometers and included some level of noise. However, the trend of VJ increase with increasing SP intensity and decreasing culture density was consistent for all fluorometers and both species (Fig. 1). The primary difference among fluorometers was related to the inability of AquaPen to adjust detector settings, which resulted in detector saturation in many conditions tested, especially when using culture suspensions with Chl *a* concentration > 0.6 mg L^−1^ (Supplementary Figures S2-S3).

#### The interplay between culture density and saturation pulse intensity

The relative FJ level depends not only on the redox states of PSII (Schansker and Strasser 2005) and the PQ pool (Tóth et al. 2007) but also on the intensity of the SP (Strasser et al. 1995; Tomek et al. 2001; Schansker et al. 2011), which is closely related to the density of the liquid suspension culture. To dissect the interplay between SP intensity and culture density, we measured OJIP transients in both *Synechocystis* and *Chlorella* cultures across a density range of 0.1 - 24.6 mg Chl *a* L^−1^, using FL 6000, Multi-Color PAM and AquaPen fluorometers with SP intensities ranging between 975 and 7 500 µmol photons m^−2^ s^−1^ (Supplementary Table S1, Supplementary Figures S2-S3).

In general, VJ was increasing with increasing SP intensity and with decreasing culture density (Fig. 1), consistently with previous works. At low light intensity, the rate of PSII excitation becomes so low that the exchange of a reduced PQ for an oxidized one is not limiting anymore, resulting in no clear J step (Tomek et al. 2001). Such results were also measured here (Supplementary Figure S3). In cell suspensions with high culture density, a significant proportion of the emitted fluorescence can be reabsorbed and/or scattered (Du et al. 2016; Kumar Panigrahi and Kumar Mishra 2019). This was also reflected in our measurements: while absolute fluorescence yield increased from low to moderate culture densities, it declined at higher cell concentrations (Supplementary Figure S2).

These results clearly demonstrate that to estimate PQ-redox from the VJ parameter reliably, the fluorescence measurement requires i) SP of consistent and appropriate intensity, high enough to secure rapid QA- accumulation and ii) consistent and low culture density, to minimize fluorescence scattering and reabsorption. The apparent shift in the VJ parameter across varying SP intensities and culture densities (Fig. 1A-D) thus reflects mainly technical limitations, rather than actual redox changes.

#### Determination of the FJ value

The exact position of FJ along the OJIP transient can be determined in multiple ways. The traditional and most common method is to detect inflection points of the OJIP fluorescence rise, with FJ defined as the point typically occurring around 2 ms in dark-acclimated samples (Strasser et al. 1995). Complementary approaches define FJ either as a local minimum in the second derivative of the fluorescence signal around 2 ms (Tomek et al. 2001; Akinyemi et al. 2023) or as a local maximum in the curvature of the fluorescence rise (Xia et al. 2019). However, these alternative methods may result in slightly miscalculated both FJ timing and amplitude. For this reason, we identified both FJ and FJ’ as inflection points occurring the closest to the typical FJ timing in dark-acclimated samples, i.e. 2 ms (see Material and Methods for details).

### VJ shift upon DCMU treatment of dark-acclimated cultures

The OJIP transient reflects the rate of forward electron transport from PSII during SP and provides information on the redox state of the PQ pool through the VJ parameter. The most straightforward way to demonstrate this is to measure the OJIP transient in the presence of DCMU (3-(3,4-dichlorophenyl)-1,1-dimethylurea). DCMU binds to the QB pocket of PSII and interrupts linear electron flow from PSII to PQ (Velthuys 1981). This effect leads to a rapid fluorescence rise along with the accumulation of QA-(Schreiber et al. 1986). When QB is displaced from its binding site, even low intensity of ML can lead to remarkable QA reduction (Tóth et al. 2005). This is due to the much slower rate of QA- recombination with the PSII donor side compared to forward electron transport from QA- to QB (Lazár 2003; Kalaji et al. 2014). Therefore, prior to measuring the DCMU-treated sample, no ML was used.

When OJIP transients were recorded in dark-acclimated *Synechocystis* and *Chlorella* cultures in the presence of 0.1 nM - 20 µM DCMU, the fluorescence rise kinetics became gradually steeper with increasing inhibitor concentrations (Schansker et al. 2011, 2014), as manifested by an increase in VJ and disappearance of the typical I and P steps along the OJIP transient (Fig. 2). At the highest tested DCMU concentrations (10-20 µM), only the O-J fluorescence phase was observed, with the fluorescence maximum reached already at the J step, in line with previous works (Tóth et al. 2005).

**Fig. 2.**
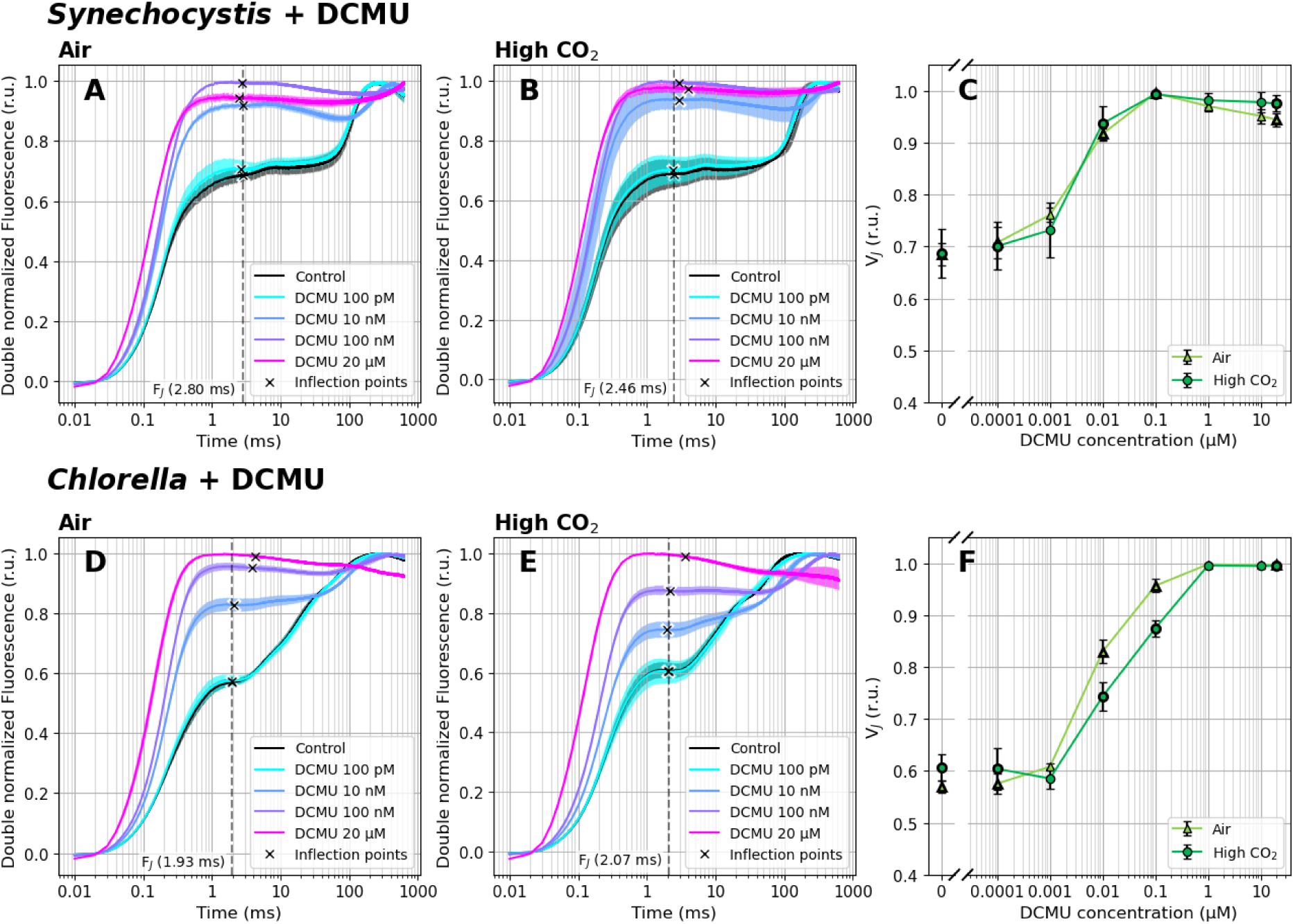
Fast Chl *a* fluorescence induction kinetics in the presence of 0 - 20 µM DCMU. Cultures of *Synechocystis* (A-C) and *Chlorella* (D-F) were pre-cultivated in a Multi-Cultivator under 25 µmol photons m^−2^ s^−1^ of white light under air (A, D) and 0.5% CO2 (B, E), and were dark-acclimated for 20 min in the presence of DCMU prior to the fluorescence measurement. After dark acclimation, the OJIP curves were recorded using a Multi-Color PAM and the parameter VJ (C, F) was calculated according to Eq. (1). OJIP curves in panels A-B and D-E were double normalized between FO and FM. The raw OJIP curves are provided in Supplementary Figure S4. FJ timing was identified as the inflection point of the fluorescence signal closest to 2 ms, and is marked by crosses in panels A-B and D-E. For simplicity, a single FJ timing (marked by the dashed line) was used for the VJ analysis within each treatment, corresponding with FJ identified in the control culture. Data represent averages ± SD, n = 4.

Since DCMU inhibits linear electron flow from PSII to PQ, the PQ pool becomes oxidized upon DCMU treatment - as measured many times previously (Kruk and Karpinski 2006; Khorobrykh et al. 2020; Virtanen and Tyystjärvi 2023). However, this PQ pool oxidation cannot be directly detected by the OJIP transient when the PSII reaction centers are closed. To measure PQ-redox changes upon DCMU treatment, simultaneous measurements of Chl *a* fluorescence and P700^+^ re-reduction kinetics would be required (Schansker et al. 2005; Belyaeva et al. 2016). The closure of PSII reaction centers in DCMU-treated cultures thus excludes the usage of the OJIP transient alone for PQ-redox estimation. We note that PQ pool oxidation over longer timescales - for instance, during state transitions - can be monitored even in the presence of DCMU (Bernát et al. 2018).

In control (i.e. untreated, dark acclimated) cultures, the VJ level was higher in *Synechocystis* compared to *Chlorella* (Fig. 1), suggesting a more reduced PQ pool in *Synechocystis*. In cyanobacteria, respiratory enzymes such as NAD(P)H dehydrogenase-like complex type 1 (NDH-1) and succinate dehydrogenase (SDH) are localized directly in the TM and transfer electrons from respiratory metabolism to TOs *via* PQ/PQH2 (Vermaas 2001). As reviewed earlier, this causes a partial PQ pool reduction in darkness (Stirbet et al. 2019). Quantitative PQ/PQH2 measurements allow for a deeper understanding of this phenomenon. After dark acclimation, a portion of reduced PQH2 within the photoactive PQ pool was found similar in *Synechocystis* sp. PCC 6803, *Chlamydomonas reinhardtii* and *Arabidopsis thaliana* (10-25%). On the other hand, the absolute size of the photoactive PQ pool was larger in *Synechocystis* than in *Chlamydomonas* or *Arabidopsis*: 47-55% *vs*. approx. 30%, respectively (Kruk and Karpinski 2006; Khorobrykh et al. 2020; Virtanen and Tyystjärvi 2023). The absolute amount of reduced PQH2 was therefore also higher in *Synechocystis* than in *Chlamydomonas* or *Arabidopsis* (30-35% *vs*. 7-15%, respectively). However, it has to be noted that the quantitative data for *Synechocystis* vary depending on the extraction procedure. Using ice-cold ethyl acetate, 25-32 PQ molecules were extracted per 1000 Chl *a* molecules, whereas petroleum ether increased this ratio approximately 1.5-2 times - which also lowered the fraction of PHQ2 reduced in darkness to about 10% (Schuurmans et al. 2014).

In addition, it has been reported that PQ-redox after dark acclimation can vary between cyanobacteria strains (Misumi et al. 2016). However, it is generally accepted that PQ pool is partially reduced in the dark in cyanobacteria - leading to relatively high FO (and/or FO’) compared to algae or plants (Campbell et al. 1998; Stirbet et al. 2019) - as confirmed also here (Supplementary Figures S4-S5). The partially reduced PQ is also assumed to be one of the factors forcing cyanobacteria to State II in the darkness (Calzadilla and Kirilovsky 2020), which is characterized by an increased absorption cross-section of PSI (σI) and decreased absorption cross-section of PSII (σII). State II ultimately leads to an underestimation of the FM value, which can only be determined correctly with maximal σII, i.e. in State I (Bhatti et al. 2021). Several methods exist to overcome these limitations and accurately determine FO and FM. For true FO determination, pre-illumination by weak far-red or blue light can be used. Such treatment enhances the activity of PSI, which leads to PQ pool oxidation and further induction of State I. However, even with such pre-treatment, FO will still be overestimated unless corrected for the contribution of fluorescence from PSI and/or free phycobilisomes (PBS) (Schuurmans et al. 2015; Ogawa and Sonoike 2016; Stirbet et al. 2019). For the correct FM determination of dark-acclimated samples, DCMU is often used since it keeps all QA reduced (Campbell and Oquist 1996).

Based on the results of the VJ parameter, reaching the maximum value of 1 under the highest DCMU concentrations ≥ 0.1 µM (*Chlorella*) or ≥ 0.01 µM (*Synechocystis*, Fig. 2), the full QA reduction and complete PSII closure (possibly involving the induction of PSII conformational changes (Sipka et al. 2021)) was confirmed also here. The increased sensitivity to DCMU in *Synechocystis* can be related to the different permeability of the inhibitor (Senger 1977). VJ levels in the presence of DCMU were largely independent of the carbon saturation level in both *Chlorella* and *Synechocystis* strains (Fig. 2). This fits the expectations: even though the CO2 level can alter acceptor side limitation of PSI (Y(NA)), DCMU makes the fluorescence response insensitive to any changes downstream of PSII. Alongside VJ, also FO values increased with higher DCMU concentrations (Supplementary Figure S4), as a result of progressive inability to transfer electrons from QA- to QB (Lazár 2003; Tóth et al. 2005).

### VJ’ dynamics during dark-to-light transition

Chl *a* fluorescence has proven effective for monitoring PQ-redox changes during dark-to-light transition. When the OJIP fluorescence rise was recorded in the absence of inhibitors, a transient VJ’ increase was found in *Chlorella* cultivated under air (Figs. 3C, F). This was likely related to the delayed activation of ferredoxin-NADP^+^-oxidoreductase (FNR) and/or Calvin-Benson-Bassham (CBB) cycle upon illumination, which prevented efficient oxidation of PQH2 by the linear electron transport under AL. FNR can be activated within a few seconds (Nikkanen et al. 2018; Kramer et al. 2021) or in a minute after a dark-to-light transition (Harbinson and Hedley 1993). Enzymes of the CBB cycle get activated during the first minutes of illumination (Michelet et al. 2013). Since CBB is ATP-dependent, formation of a pH gradient across the TM is essential to support CBB operation at a full rate. Previous observations on pH changes in chloroplasts during dark-to-light transitions revealed a stable pH after ∼1-2 minutes of illumination both in the thylakoid lumen and the cytoplasm (Heldt et al. 1973). This is roughly consistent with the time of VJ’ peak observed in air-cultivated *Chlorella* here (Fig. 3F). After 5 min of illumination, when FNR and CBB get fully activated and NPQ and state transitions can be expected to form a quasi-steady-state, the VJ’ value stabilized, reflecting a more reduced PQ pool in light-acclimated cultures, compared to dark-acclimated state (Khorobrykh et al. 2020; Virtanen and Tyystjärvi 2023). We note that the timing of VJ’ increase, decline and stabilization coincides with the ‘slow’ Chl *a* fluorescence kinetics measured during dark-to-light transition (Lichtenthaler et al. 2005; Kalaji et al. 2014).

**Fig. 3.**
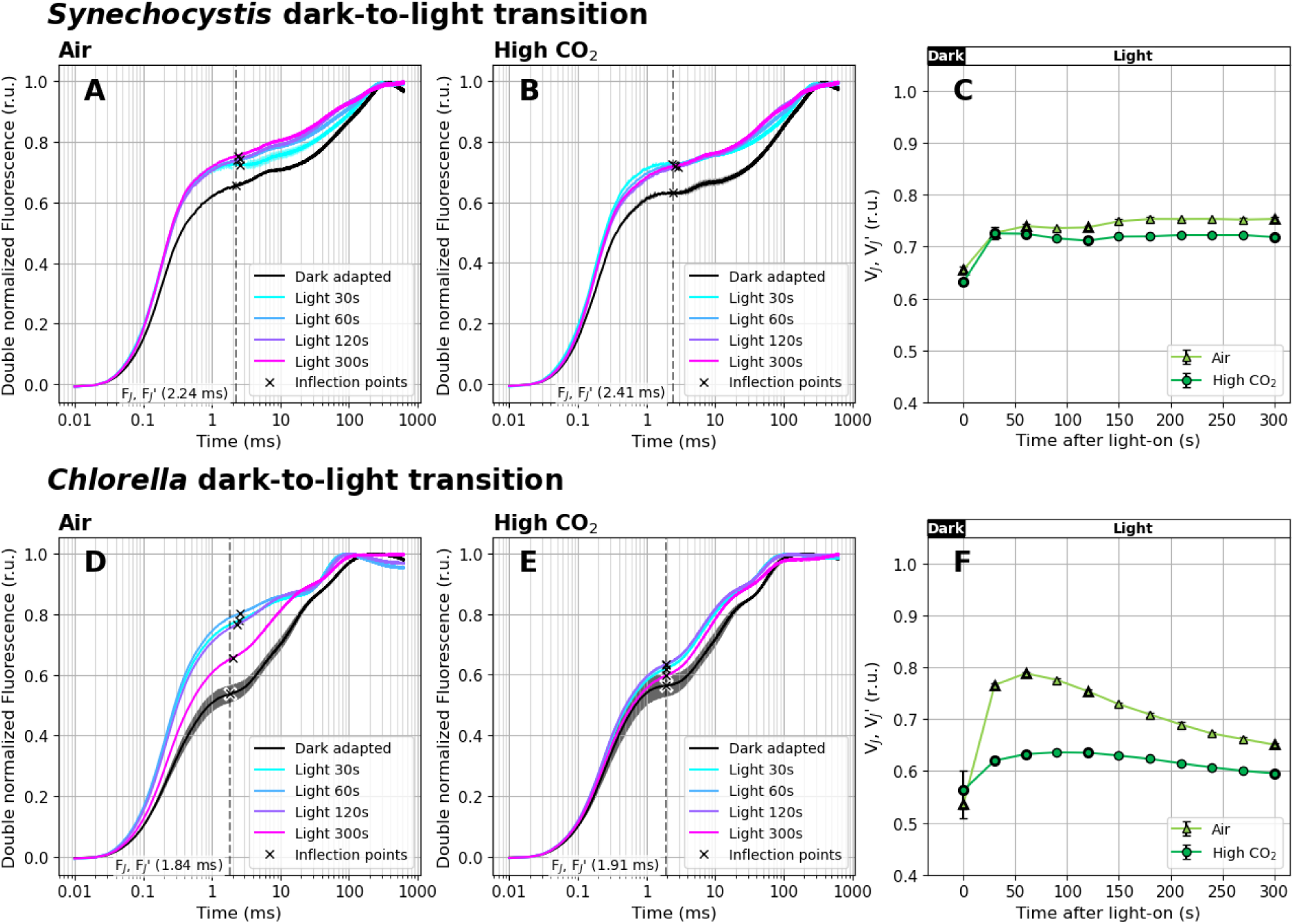
Fast Chl *a* fluorescence induction kinetics during dark-to-light transition (100 µmol photons m^−2^ s^−1^ of white AL). Cultures of *Synechocystis* (A-C) and *Chlorella* (D-F) were pre-cultivated in a Multi-Cultivator under 25 µmol photons m^−2^ s^−1^ of white light under air (A, D) and 0.5% CO2 (B, E) and were dark-acclimated for 20 min prior to the fluorescence measurement. After dark acclimation, the OJIP curves were recorded using Multi-Color PAM and the parameters VJ and VJ’ (C, F) were calculated according to Eq. (1) and Eq. (2), respectively. The OJIP curves in panels A-B and D-E were double normalized between FO and FM or between FO’ and FM’(dark- and light-acclimated cultures, respectively). The raw OJIP curves are provided in Supplementary Figure S5. FJ and FJ’ timing, identified as the inflection points of the fluorescence signal closest to 2 ms, are marked by crosses in panels A-B and D-E. For simplicity, a single FJ’ timing was used for VJ’ analysis (marked by the dashed line), based on FJ identified in the dark-acclimated culture. Data represent averages ± SD, n = 4.

In *Chlorella* cultivated under high CO2, VJ’ was more stable compared to air-acclimated cultures. Higher CO2 levels likely resulted in a faster activation of the CBB cycle enzymes. Under high CO2, a greater portion of RuBisCO is located in the chloroplast stroma, whereas under carbon limitation, key CBB cycle enzymes are found in the pyrenoid, an organelle of carbon-concentrating mechanism (CCM) (He et al. 2023). The CCM secures CO2 accumulation in the pyrenoid by introducing several diffusion barriers that prevent CO2 leakage back to the stroma (Fei et al. 2022). As a result, in high CO2 conditions, more substrate becomes readily available to RuBisCO which likely can be expected to minimize FNR and CBB activation times, and to mitigate Y(NA) at the onset of illumination - in good agreement with the VJ’ dynamics measured here (Fig. 3F).

In *Synechocystis*, the surprisingly stable VJ’ during 5 min of illumination was likely a result of the activity of terminal oxidases present in the TM - cytochrome *bd* quinol oxidase (Cyd) and cytochrome *c* oxidase (COX) (Howitt and Vermaas 1998; Berry et al. 2002; Lea-Smith et al. 2013). These enzymes can take up electrons from PQH2, independently of the activation status of FNR and the CBB cycle, and therefore prevent PQ pool over-reduction at the onset of illumination. We note, however, that measurements with mutants lacking these TOs would be needed to provide direct evidence for this hypothesis. *Synechocystis* can further balance PQ-redox through PBS decoupling and/or state transitions (Stirbet et al. 2019). Since FO’ did not decrease in our measurements, PBS decoupling was unlikely under 100 µmol photons m^−2^ s^−1^ (Supplementary Figure S5). On the other hand, relatively high amount of blue photons in the white AL spectrum (Supplementary Figure S1) likely contributed to a partial State I transition under illumination (Calzadilla and Kirilovsky 2020), resulting in FM’ increase (Supplementary Figure S5).

Besides the shifts in VJ’, the dark-to-light transition caused shifts in the timing of FP (“Peak” of the OJIP curve). In the light, FP’ appeared earlier compared to dark in both strains: in *Chlorella* under low and high CO2, FP was identified at 115±17 ms and 86±13 ms in the dark and at 62±1 ms and 68±0 ms after 30 s at light, respectively. In *Synechocystis*, FP occurred at 333±12 ms and 340±27 ms in the dark and at 245±3 ms and 236±6 ms after 30 s at light, under low and high CO2, respectively. The earlier FP’ appearance at light persisted in *Chlorella*, while in *Synechocystis*, FP’ gradually shifted to a later time over the 5-minute illumination period (Supplementary Figure S5). Earlier appearance of FP’ can be related to an enhanced Y(NA) (Stirbet and Govindjee 2012), and/or the activation of the xanthophyll cycle and NPQ in green algae and plants (Short et al. 2023). Upon Y(NA), electrons from PETC cannot be taken up by PSI and donated to downstream carriers, which enhances the fluorescence yield of PSII. During xanthophyll cycle activation, the pH gradient across the TM gets smaller, which allows Cyt *b*6*f* to transport electrons more efficiently (Gog et al. 2019). Further, it has been shown that the IP phase gets suppressed upon activation of the CBB cycle, leading to earlier appearance of the apparent FP’ (Schansker et al. 2006). Contrary, FP’ can be delayed by an increased pH gradient across the TM, which can downregulate the activity of Cyt *b*6/*f* (Joliot and Johnson 2011), and/or State I→State II transition upon illumination, reducing the amount of light reaching PSII (Tomek et al. 2001).

Indeed, actinic illumination led to QA- accumulation within PSII. In dark-acclimated cells, nearly all PSII reaction centers are in the open state, QA is fully oxidized, and the oxygen-evolving complex (OEC) is in the dark-stable S1 (75%) and S0 (25%) states (Jablonsky and Lazar 2008). Upon illumination by AL of moderate intensity (Figs. 3-6), a fraction of P680 (the primary electron donor of PSII) gets excited (P680*). After charge separation, this leads to a partial reduction of pheophytin (Pheo^−^) and QA (QA-) which further reduces QB and forms a new redox equilibrium with PQ/PQH2, depending on the redox state of downstream PETC components. The OJIP transient under illumination thus reflects a combined effect of light-driven QA- accumulation in PSII, and the activity of PETC components downstream of PSII, including accumulation of PQH2 within the photoactive PQ pool (Khorobrykh et al. 2020; Virtanen and Tyystjärvi 2023).

As a control to the dark-to-light transition experiment, an opposite light-to-dark shift was also performed. According to the expectations, VJ’ parameter decreased after the transition from light (10 µmol photons m^−2^ s^−1^) to dark, and the decline was more pronounced in *Chlorella* than in *Synechocystis*. This suggests gradual oxidation of the PQ pool in darkness, in agreement with previous works (Ivanov et al. 2007). It also suggests a tight control of PQ-redox in *Synechocystis* (Schuurmans et al. 2014). Full description of the dark-to-light transition experiment is provided in Supplementary material, and the results are summarized in Supplementary Figure S6. Taken together, these results confirm that the VJ and VJ’ parameters derived from the OJIP curves are sensitive enough to reflect, on a semi-quantitative scale, changes in the redox state of the PQ pool in both dark- and light-acclimated green algae and cyanobacteria cultures.

### VJ’ dynamics during dark-to-light transition in the presence of methyl viologen

To verify that the transient increase in VJ’ upon illumination in *Chlorella* was associated with FNR and the CBB cycle activation, OJIP transients during dark-to-light transition were additionally recorded in the presence of methyl viologen. This substance efficiently accepts electrons from PSI and FNR and transfers them to molecular oxygen (Fuerst and Norman 1991; Sétif 2015), competing with FNR and the CBB cycle. It therefore prevents the reduction of NADP^+^ and cyclic electron flow around PSI (PSI-CEF) (Yu et al. 1993). MV present in 1 mM concentration, after 20 min pre-incubation, prevented the transient VJ’ increase (Figs. 4C, F) observed in control samples (Fig. 3F). At a lower MV concentration (0.25 mM), the VJ’ shift under illumination was more pronounced than in the absence of MV, but less pronounced than at 1 mM MV concentration (Supplementary Figures S7-S8).

**Fig. 4.**
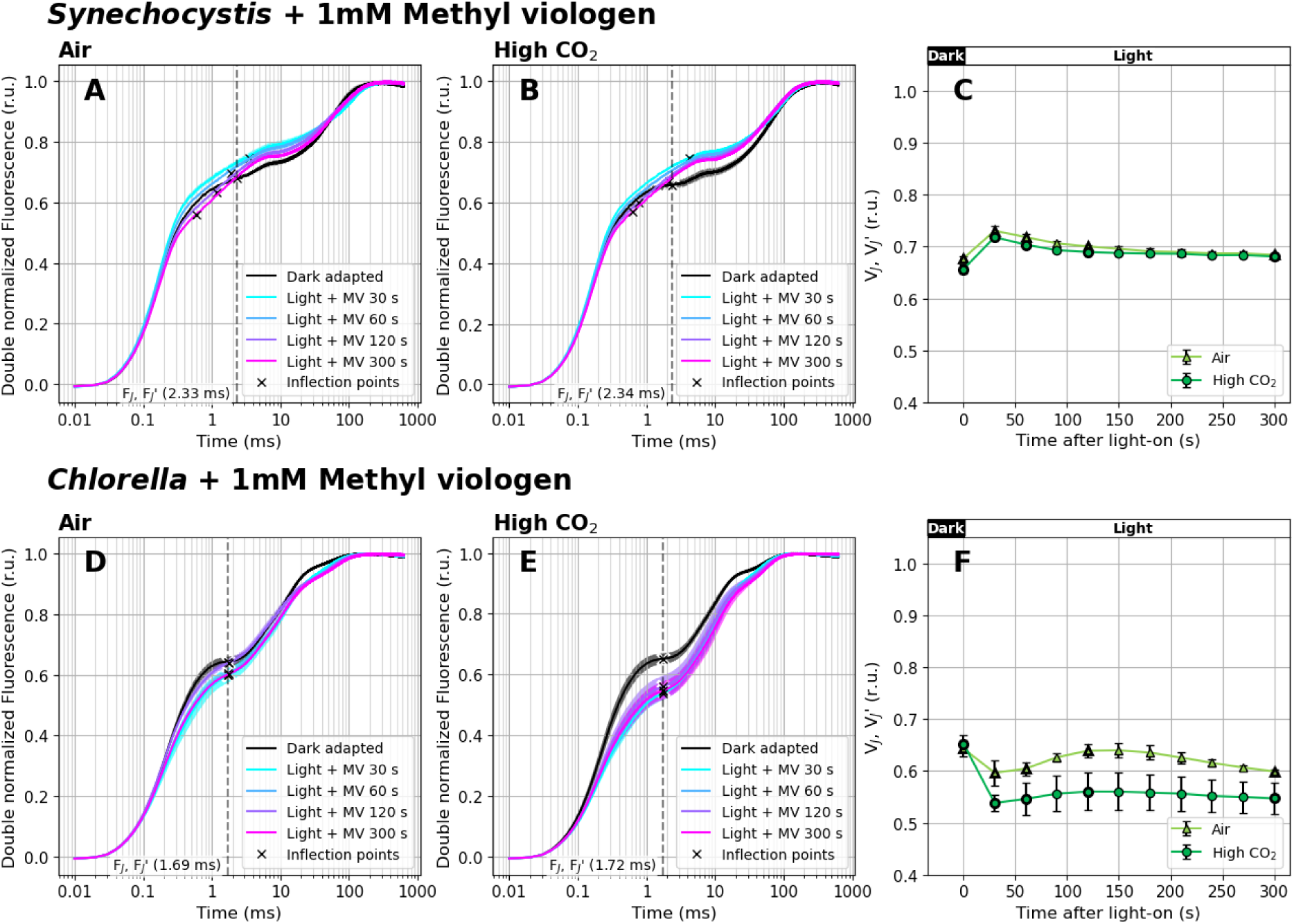
Fast Chl *a* fluorescence induction kinetics during dark-to-light transition (100 µmol photons m^−2^ s^−1^ of white light) in the presence of 1 mM methyl viologen (MV). Cultures of *Synechocystis* (A-C) and *Chlorella* (D-F) were pre-cultivated in a Multi-Cultivator under 25 µmol photons m^−2^ s^−1^ of white light under air (A, D) and 0.5% CO2 (B, E) and were dark-acclimated for 20 min prior to the fluorescence measurement in the presence of MV. After dark acclimation, OJIP curves were recorded using Multi-Color PAM and the parameters VJ and VJ’ (C, F) were calculated according to Eq. (1) and Eq. (2), respectively. OJIP curves in panels A-B and D-E were double normalized between FO and FM. The raw OJIP curves are provided in Supplementary Figure S11. OJIP curves recorded under 0.25 mM MV are provided in Supplementary Figures S7-S8. FJ and FJ’ timing, identified as the inflection points of the fluorescence signal closest to 2 ms, are marked by crosses in panels A-B and D-E. For simplicity, a single FJ’ timing was used for VJ’ analysis (marked by the dashed line), based on FJ identified in the control culture. Data represent averages ± SD, n = 4.

Interestingly, in dark-acclimated cultures, the presence of MV led to VJ increase, likely due to a reduced IP phase associated with the FNR bypass (Fig. 4, Supplementary Figure S9). This shows a limited capacity of the VJ parameter to efficiently reflect PQ-redox in samples with distinct Y(NA) states. Opposite to previously reported MV treatment in vascular plants (Schansker et al. 2005), we observed a relative increase in the overall OJIP amplitude in both *Chlorella* and *Synechocystis* cultures (compared to the MV-free control, Supplementary Figure S10). Previous studies have shown that when the FNR pathway is fully bypassed in vascular plants, the IP phase entirely disappears (Schansker et al. 2005). In our measurements the IP phase remained detectable in the presence of MV (Supplementary Figures S7-S8, S11), albeit reduced compared to control cultures (Supplementary Figures S9). The IP phase in *Chlorella* was smaller than reported previously in vascular plants (Schansker et al. 2005), and in *Synechocystis* it was barely visible. These results show that a direct comparison of OJIP trends for vascular plants, algae and cyanobacteria requires precaution. Nevertheless, the 20 min pre-incubation with MV was still long enough to partially prevent Y(NA) at the onset of illumination. Increasing the sink for PSET electrons downstream of PSI is known to promote PQ pool oxidation (Schansker et al. 2005; Khorobrykh et al. 2020). Lower VJ’ values following MV treatment, relative to control (Figs. 3-4), thus confirm that the VJ’ parameter is sensitive enough to reflect transient changes in the PQ-redox even under actinic light.

### VJ’ dynamics upon the addition of glycolaldehyde under illumination

To provide additional evidence that the VJ’ parameter can reflect changes in PQ-redox under illumination, OJIP curves were further recorded under actinic light in the presence of 25 mM glycolaldehyde (GA). This compound inhibits enzymes of the CBB cycle (Stokes and Walker 1972), and its presence can thus be expected to induce Y(NA) and to increase VJ and VJ’ due to PQ pool reduction. Our results confirmed this assumption: when GA was added to the light-acclimated cultures, it led to VJ’ increase already after 30 s of exposure (Fig. 5, Supplementary Figure S11).

**Fig. 5.**
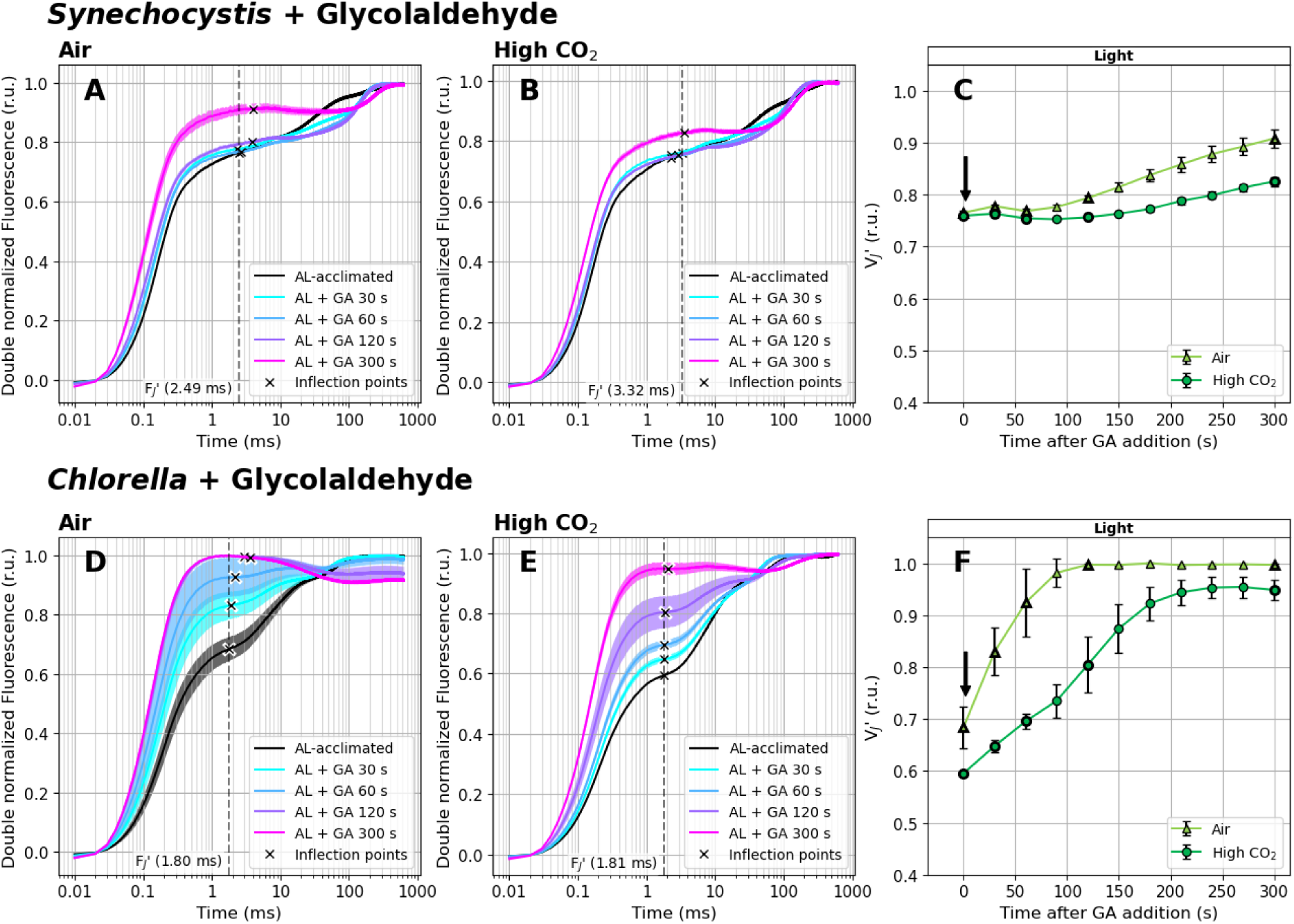
Fast Chl *a* fluorescence induction kinetics measured under white actinic light (100 µmol photons m^−2^ s^−1^) in the presence of 25 mM glycolaldehyde (GA). Cultures of *Synechocystis* (A-C) and *Chlorella* (D-F) were pre-cultivated in a Multi-Cultivator under 25 µmol photons m^−2^ s^−1^ of white light under air (A, D) and 0.5% CO2 (B, E) and were further light-acclimated in the Multi-Color PAM for 10 min prior to the fluorescence measurement under 100 µmol photons m^−2^ s^−1^ of white actinic light. After light acclimation, OJIP curves were recorded using Multi-Color PAM and the parameter VJ’ (C, F) was calculated according to Eq. (2). Timing of GA addition, right after the first OJIP measurement, is marked by arrows in panels C and F. OJIP curves in panels A-B and D-E were double normalized between FO’ and FM’. The raw OJIP curves are provided in Supplementary Figure S12. FJ’ timing, identified as the inflection point of the fluorescence signal closest to 2 ms, is marked by crosses in panels A-B and D-E. For simplicity, a single FJ’ timing (marked by the dashed line) was used for the VJ’ analysis within each treatment, corresponding with FJ’ identified in the control cultures. Data represent averages ± SD, n = 4.

**Fig. 6.**
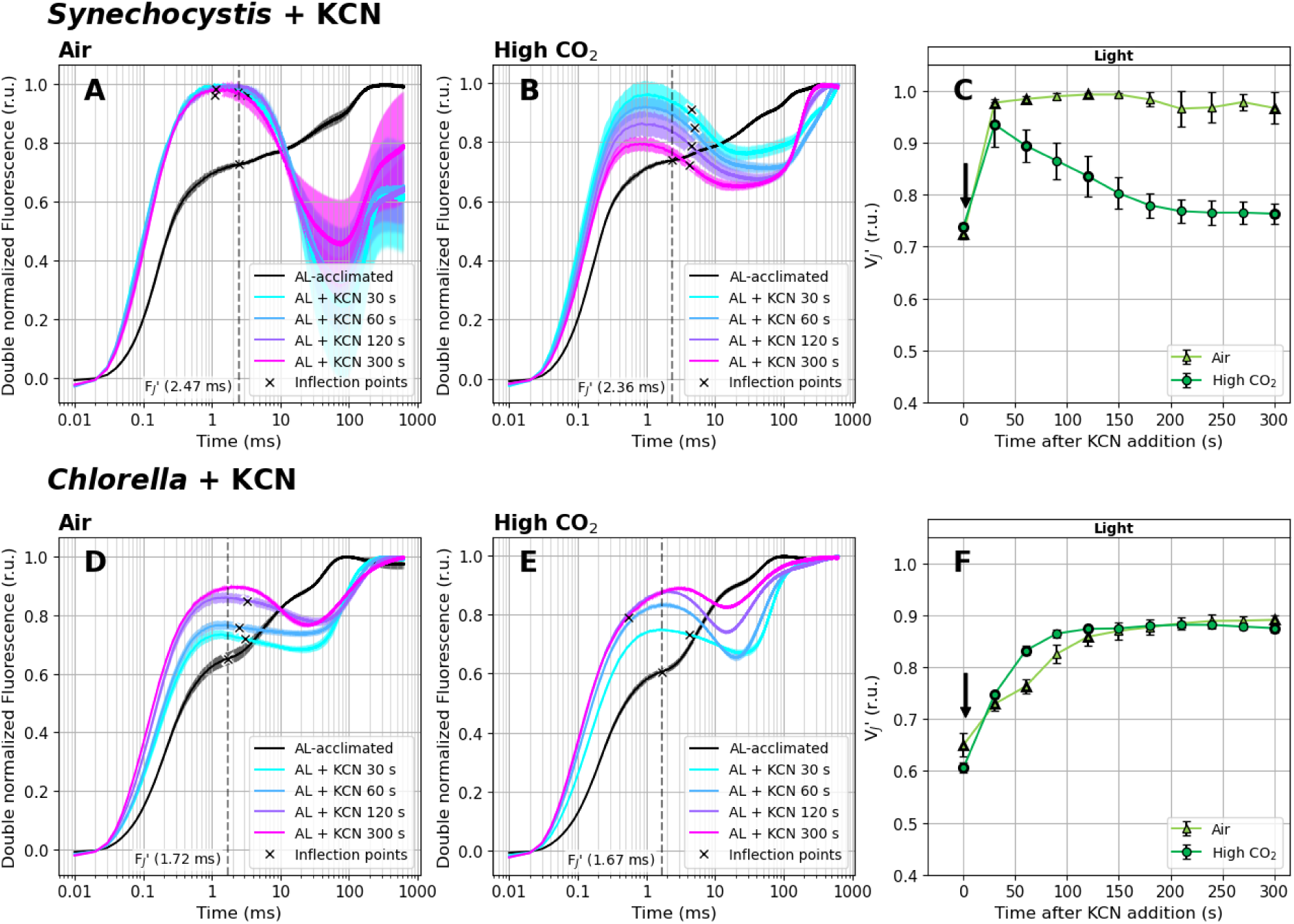
Fast Chl *a* fluorescence induction kinetics measured under white actinic light (100 µmol photons m^−2^ s^−1^) in the presence of 1 mM potassium cyanide (KCN). Cultures of *Synechocystis* (A-C) and *Chlorella* (D-F) were pre-cultivated in a Multi-Cultivator under 25 µmol photons m^−2^ s^−1^ of white light under air (A, D) and 0.5% CO2 (B, E) and were further light-acclimated in the Multi-Color PAM for 10 min prior to the fluorescence measurement under 100 µmol photons m^−2^ s^−1^ of white actinic light. After light acclimation, OJIP curves were recorded using Multi-Color PAM and the parameter VJ’ (C, F) was calculated according to Eq. (2). Timing of KCN addition, performed immediately after the first OJIP measurement, is marked by arrows in panels C and F. OJIP curves in panels A-B and D-E were double normalized between FO’ and FM’. The raw OJIP curves are provided in Supplementary Figure S13. FJ’ timing, identified as the inflection point of the fluorescence signal closest to 2 ms, is marked by crosses in panels A-B and D-E. For simplicity, a single FJ’ timing (marked by the dashed line) was used for the VJ’ analysis within each treatment, corresponding with FJ’ identified in the controlled culture. Data represent averages ± SD, n = 4.

The initial value of VJ’, reflecting PQ-redox after 10 min of acclimation to AL of PFD of 100 µmol photons m^−2^ s^−1^, was the highest in *Synechocystis*, lower in *Chlorella* cultivated under air and the lowest in *Chlorella* under high CO2 (Figs. 5C, F) in accordance with other measurements (Figs. 3, 6). The VJ’ shift upon GA addition was dependent on the CO2 level under which both strains were cultivated. In air-cultivated cultures, VJ’ was increasing at a higher rate and reached higher levels compared to cultures cultivated under high CO2 (Figs. 5C, F). In *Chlorella*, VJ’ reached the maximum possible value of 1 at about 120 s after GA addition, reflecting a highly reduced PQ pool and a lack of alternative routes of PQH2 oxidation. In *Chlorella* cultivated under high CO2, the VJ’ rise was slower and it stabilized around the value of 0.95, suggesting also a highly reduced yet slightly more oxidized PQ pool compared to low CO2 conditions - in agreement with previous works (Schuurmans et al. 2014; Khorobrykh et al. 2020). The difference in PQ-redox after GA treatment between high- and low-CO2 acclimated cultures can be attributed simply to higher CBB cycle substrate being available, and therefore higher GA concentrations needed to fully inhibit the activity of the CBB cycle enzymes under high CO2 conditions. In *Synechocystis*, VJ’ shifts were less pronounced compared to *Chlorella* (Fig. 5), again, likely due to the presence of TOs at TM, balancing PQ reduction upon GA addition (see next section for details).

GA also triggered an increase in FO’ values and changes in FP’ timing (Supplementary Figure S12). The FO’ increase was likely a result of redox back-pressure on QA, analogous to the effect of DCMU treatment (Supplementary Figure S4). However, with GA treatment this back-pressure can be expected to originate outside of PSII, in particular in strong Y(NA) that further creates back-pressure on PQH2. In *Synechocystis*, the FO’ increase was lower compared to *Chlorella*, again likely due to the presence of respiratory enzymes at TM that balanced PQ-redox upon GA application. After 300 s of incubation in the presence of GA, the appearance of FP’ was slightly delayed compared to control in both strains (Supplementary Figure S12).

### VJ’ dynamics upon addition of KCN under illumination

To confirm that terminal oxidases balance PQ-redox in *Synechocystis*, OJIP transients were further recorded under actinic light in the presence of 1 mM KCN. This substance inhibits TOs such as cytochrome c oxidase (COX) and cytochrome *bd* quinol oxidase (Cyd), both present in *Synechocystis*, with inhibition constant KI as low as 7 and 27 µM, respectively (Pils and Schmetterer 2001). However, KCN also has several side effects, such as inhibition of the activity of plastocyanin and therefore disruption of linear electron flow between PSII and PSI (Redinbo et al. 1994). It may also disrupt the activity of PSII or CBB cycle (Sanakis et al. 1994; Tjus et al. 1998; Hill et al. 2014), and cause a decrease of pH gradient across the TM (Miller et al. 2021). Despite such KCN non-specificity, the terminal oxidases are by far the most sensitive KCN targets (Pils and Schmetterer 2001; Hill et al. 2014; Santos-Merino et al. 2021; Kusama et al. 2022).

Here, KCN was added to light-acclimated cultures and its effect on the VJ’ parameter was evaluated for 5 min right after its addition (in 30 s intervals). This setup prevented the above-described undesired KCN effects on PSII, CBB, plastocyanin or other PETC components during pre-incubation. Similar to Figs. 3 and 5, the VJ’ values prior to KCN addition, i.e. after 10 min of acclimation under 100 µmol photons m^−2^ s^−1^, were the highest in *Synechocystis*, lower in *Chlorella* under air and the lowest in *Chlorella* under high CO2 (Figs. 6C, F). KCN addition induced VJ’ increase in both species, and this increase was much faster and more pronounced in *Synechocystis*, where the VJ’ reached values of 0.94±0.04 and 0.98±0.01 (high and low CO2, respectively) already 30 s after KCN addition. In *Chlorella*, VJ’ was rising for about 2 min before reaching a stable value of around 0.88±0.01 (Fig. 6).

The high VJ’ values after KCN addition, reflecting strong reduction of the PQ pool, are in full agreement with previous results (Fukunaga et al. 2024). The fast VJ’ increase in *Synechocystis* upon KCN addition further confirmed the role of TOs in PQ-redox balancing. Under AL, PQ is reduced by PSII as well as through metabolism *via* SDH and NDH-1, and oxidized by TOs and PSI. Inactivating TOs thus creates redox imbalance, which is higher in *Synechocystis* compared to *Chlorella* in which PSET and respiratory electron flow enzymes are located in different organelles. In *Synechocystis* cultivated under air, VJ’ remained high (0.97±0.03), whereas under high CO2, the initial rise was only transient and VJ’ decreased to 0.76±0.02 after 5 min of illumination (Fig. 6). In *Synechocystis*, the absolute amounts of the CBB cycle enzymes and/or terminal oxidases was previously found independent of CO2 level (Jahn et al. 2018; Mustila et al. 2021). Therefore, the slightly lower and over time decreasing VJ’ in *Synechocystis* cultivated under high CO2 was likely a consequence of either more efficient CO2 fixation resulting from higher carbon availability - which partially decreases Y(NA) and leads to a more oxidized PQ pool compared to air conditions (Schuurmans et al. 2014; Khorobrykh et al. 2020) - or of less pronounced electron flow *via* SDH and NDH-1.

The VJ’ increase in *Synechocystis* was accompanied by FO’ rise (Supplementary Figure S13). This is not surprising, as due to the inhibition of TOs by KCN, the ability of PQ to accept electrons from QA- is significantly reduced. This back-pressure on QA through PQH2 leads again to QA- accumulation, analogous to the effect of GA observed in *Chlorella* (Supplementary Figure S12). However, in the case of KCN, extra back-pressure on QA was likely induced by KCN binding to the non-heme iron of PSII (Sanakis et al. 1994), impairing the electron transport between QA and QB similar to DCMU (Vermaas et al. 1994). We note, however, that PSII inhibition was likely still only partial after 5 min of KCN treatment under AL. In addition, the FO’ value was increasing over the 5 min-period in *Chlorella*, whereas it started to decline after 2 min in *Synechocystis*. This was likely related to a partially induced State I → State II transition (Calzadilla and Kirilovsky 2020), as reflected by FM’ decrease (Supplementary Figure S13). The FO’ rise can be also related to the accumulation of reactive oxygen species (ROS) upon partial inactivation of the CBB cycle by KCN (Siegień and Bogatek 2006).

Besides a shift in FO’, a transient dip in the fluorescence induction was also recorded between FJ’ and FP’ upon KCN addition in *Synechocystis* (Figs. 6A-B). This was likely a consequence of lower affinity of KCN to the enzymes of CBB cycle than to TOs: after impairing TOs, the FJ’ fluorescence yield increased, however, unlike in the presence of DCMU (Fig. 2) electrons were still able to flow from PQ toward PSI (Schansker et al. 2005), as CBB was not blocked by KCN as strongly as the TOs (Hill et al. 2014).

In *Chlorella*, VJ’ was increasing at a slower rate, and reached lower level compared to *Synechocystis* (Fig. 6). The only known terminal oxidase present in the chloroplast of *Chlorella* is plastid terminal oxidase (PTOX), which is, however, resistant to KCN (Fu et al. 2009). Therefore, the slower VJ’ increase in *Chlorella* upon KCN addition, compared to *Synechocystis*, further proves that KCN did not inhibit chloroplastic TOs in *Chlorella*. Indeed, in *Chlorella*, KCN likely inhibited TOs and ATP production in mitochondria, and reduced NAD(P)H export from the chloroplast to mitochondria *via* metabolic shuttles (Noguchi and Yoshida 2008). However, in contrast to *Synechocystis*, where the enzymes of both PSET and respiratory electron flow are localized in the same TM, in *Chlorella* these processes are connected indirectly. The VJ’ increase can be further linked to gradual inhibition of plastocyanin and the CBB cycle enzymes (Wishnick and Lane 1969; Berg and Krogmann 1975), both contributing to redox back-pressure on PQ. KCN also inhibits carbonic anhydrase (Lindahl et al. 1993), which can further reduce CBB activity.

These effects likely resulted in strong PQ reduction in *Chlorella* after 5 min of KCN treatment, as reflected in high VJ’ values (Fig. 6F). Additionally, a dip in the fluorescence signal between FJ’ and FP’ was observed (Figs. 6D-E). Consistent with the results in *Synechocystis* (Fig. 6C) and previous observations under anaerobic conditions in vascular plants (Tóth et al. 2007), this dip further indicates a highly reduced PQ pool.

### VJ’ shift under high light

Results presented in the previous sections show that the VJ and VJ’ parameters reliably reflect PQ-redox shifts in darkness and under moderate light intensity. To provide additional information on the relationship between PSII closure and the ability of the VJ parameter to reflect PQ-redox, we recorded OJIP curves during transition from low (25 µmol photons m^−2^ s^−1^) to high light (HL, 1 500 µmol photons m^−2^ s^−1^).

Upon HL treatment, the excitation rate of P680 increases significantly, leading to higher rate of charge separation within PSII. The reaction center of PSII, P680, is reduced on the donor side by the redox-active tyrosine residue (Tyr-Z) of the D1 protein, which receives an electron from the oxygen-evolving complex (OEC). At the acceptor side of PSII, the electrons are transferred through pheophytin to QA and QB. Since the electron transport between QA and QB is several orders of magnitude faster than PQ reduction to PQH2 (Lazár 2003), QA- accumulates rapidly upon HL exposure. The OJIP transient, measured in HL-acclimated cultures, is therefore strongly affected by the pre-accumulated QA-. This limits the ability of the fluorescence signal to reflect changes downstream of PSII, including PQ-redox shifts. After the transition from low to high light, the amount of PQH2 within the total PQ pool increased previously from ∼30-50% to ∼75% in *Synechocystis (Khorobrykh et al. 2020)*. In addition, the size of the photoactive pool increased under high light (Pralon et al. 2019). Here, the VJ’ parameter increased only from 0.72±0.03 to 0.8±0.01 in *Synechocystis*, and from 0.53-0.56 to 0.73-0.81 in *Chlorella* (pooled between CO2 treatments; Figs. 7C, F). This clearly demonstrates that under HL, the range of the VJ’ parameter reflecting PQ-redox shifts narrows down - limiting this parameter to reflect PQ-redox under high light treatment.

**Fig. 7.**
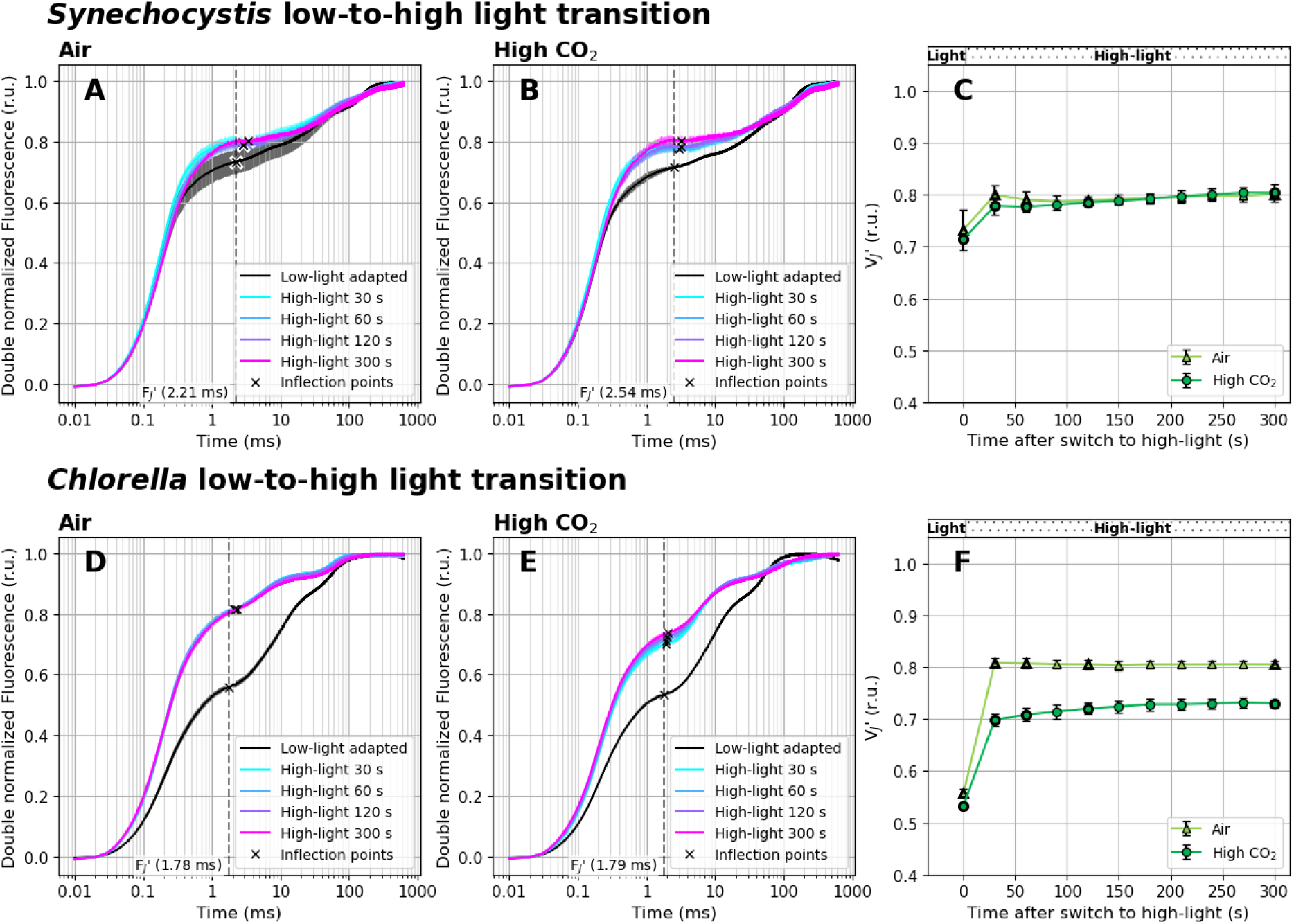
Fast Chl *a* fluorescence induction kinetics during high light treatment. Cultures of *Synechocystis* (A-C) and *Chlorella* (D-F) were pre-cultivated in a Multi-Cultivator under 25 µmol photons m^−2^ s^−1^ of white light under air (A, D) and 0.5% CO2 (B, E). Culture aliquotes were transferred to Multi-Color PAM and further acclimated under 25 µmol photons m^−2^ s^−1^ of white light for 5 min. After acclimation, OJIP curves were recorded, the PFD of white actinic light was increased to 1 500 µmol photons m^−2^ s^−1^, and OJIP curves were further recorded in 30 second intervals for next 5 min. The parameter VJ’ (C, F) was calculated according to Eq. (2). OJIP curves in panels A-B and D-E were double normalized between FO’ and FM’. The raw OJIP curves are provided in Supplementary Figure S14. FJ’ timing, identified as the inflection point of the fluorescence signal closest to 2 ms, is marked by crosses in panels A-B and D-E. For simplicity, a single FJ’ timing (marked by the dashed line) was used for the VJ’ analysis within each treatment, corresponding with FJ’ identified in the controlled culture. Data represent averages ± SD, n = 4.

Nevertheless, fluorescence transients upon HL treatment, including shifts in the VJ’ parameter, differed significantly in *Synechocystis* and *Chlorella* - reflecting distinct HL acclimation strategies in both strains. In *Synechocystis*, the VJ’ increase under HL, compared to low light, was less pronounced compared to *Chlorella* (Fig. 7) - suggesting a stronger PQ-redox control (Schuurmans et al. 2014). This can be again related to the activity of TOs (Lea-Smith et al. 2013), but also to functional uncoupling of PBS from PSII (Tamary et al. 2012) or to non-photochemical quenching mediated through orange carotenoid protein (OCP-NPQ; (Kirilovsky and Kerfeld 2013)). Both latter effects can lead to FM’ decrease under HL in *Synechocystis* (Supplementary Figure S14); the PBS decoupling in *Synechocystis* was measured here directly (Supplementary Figure S15). Indeed, also decoupled PBS emits fluorescence that might contribute to the overall fluorescence signal, potentially increasing FO’ (Remelli and Santabarbara 2018). We note, however, that in *Synechocystis* both PBS decoupling and OCP-NPQ likely dissipated only part of the excessive energy that this strain received under HL (Tian et al. 2011; Pfennig et al. 2024).

Additionally, redox homeostasis can be further balanced by alternative electron flows such as the water-water cycle, cyclic electron flow around PSI (Miyake 2010) or NPQ securing heat dissipation; in *Synechocystis* through OCP-NPQ and in *Chlorella* through the xanthophyll cycle. These protective mechanisms can also lead to a decline of maximal fluorescence value, FM’, as detected in both strains under HL (Supplementary Figure S14). The FM’ reduction can be further related to gradual PSII damage (Kanervo et al. 1993; Wilson et al. 2006), as prolonged limitation of forward electron transport and increased P * lifetime leads to formation of ROS both on donor and acceptor sides of PSII (Vass 2012).

## Discussion

### Advantages of the fluorescence method for PQ-redox estimation

This study evaluates the ability of OJIP fluorescence transients to reflect redox state of the PQ pool in green algae and cyanobacteria. PQ pool is a central component in the acclimation of phototrophic cells to diverse environmental conditions, and determination of its redox state is often highly desirable. Under well-defined experimental conditions, this fluorescence method can non-invasively and reliably reflect shifts in the redox state of the PQ pool *in vivo*, as long as the key assumptions and limitations are acknowledged (see below). In plant and crop research, it has also been suggested for *in situ* applications (Tóth et al. 2007).

A clear advantage of this fluorescence method is its simplicity, as it avoids PQ/PQH2 extraction (Kruk and Karpinski 2006; Schuurmans et al. 2014; Khorobrykh et al. 2020). The method requires ‘only’ a kinetic fluorometer with high light intensity and high temporal resolution (10^−6^ s, or higher). Such fluorometers are commercially available in many versions and price levels and can even be built using open-source schemes (Bates et al. 2019).

The fast Chl *a* fluorescence induction (OJIP) transient is typically measured in a dark-acclimated state, in order to reach a stable, relaxed state of the PETC. This includes relaxation of P680, quasi full oxidation of QA and QB, oxidation of the PQ pool to the maximum extent possible under a given condition, shifting OEC to a dark-stable S1 (and S0) state and inactivation of FNR and CBB-cycle. In this work, we show that the OJIP transient can reflect PQ-redox even in a light-acclimated state. Measuring pre-illuminated cultures, however, prevents correct determination of the true FO and FM values that are essential for calculating a variety of parameters of the so-called JIP test, including VJ (Stirbet et al. 2018; Tsimilli-Michael 2020). Obtaining true FO and FM is especially challenging in cyanobacteria, where FM is typically measured in the presence of DCMU, while reliable estimation of FO requires i) a full induction of State I and ii) a correction of the fluorescence signal for contributions from PSI and PBS emission (Ogawa and Sonoike 2016; Stirbet et al. 2019). Nevertheless, as we show here, the true FO and FM values are not required to reflect relative changes in the redox state of the PQ pool, as the parameter VJ’ reflects PQ-redox shifts well even in light-acclimated cultures.

### Limitations of the method

However, estimating the PQ-redox based on the VJ and VJ’ parameters has several ultimate limits. First, it becomes unsuitable when the electron transport between QB and PQ gets impaired, for instance under DCMU treatment (Fig. 2), or when the redox state of PSII substantially differs between measurements - such as during transition from low to high light (Fig. 7). The OJIP measurements in the latter case show that the more QA- accumulates in PSII prior to the OJIP measurement, the less QA remains available to form equilibrium with PETC components downstream of PSII, including PQ/PQH2. Second, the dynamic range of the VJ’ parameter (calculated using FO’ and FM’, Eq. 2) can be substantially narrower than that of VJ (based on FO and FM, Eq. 1), which may, for example, weaken correlation with PQ-redox changes between dark- and light-acclimated samples. Third, the working range of VJ and VJ’ will always reflect only the photoactive fraction of the PQ pool, rather than total PQ/PQH2 pools. The photoactive pool was estimated between 25-40% of total PQ in plants and algae and between 41-55% in *Synechocystis* previously (Kruk and Karpinski 2006; Yoshida et al. 2010; Khorobrykh et al. 2020; Mattila et al. 2020; Virtanen and Tyystjärvi 2023).

The VJ’ values obtained here can be compared with quantitative PQ/PQH2 data, measured previously under a variety of conditions and strains. In *Synechocystis* cultivated under high CO2, the amount of PQH2 within the photoactive pool increased 1.06- or 1.17-fold (depending on normalization) after the transition from dark to light at 40 µmol photons m^−2^ s^−1^ (Khorobrykh et al. 2020). Here, VJ’ increased 1.14-fold after the transition from dark to 100 µmol photons m^−2^ s^−1^. Under air, VJ’ increased 1.15-fold (Fig. 3), whereas PQH2 increased 2-fold in the previous study. In other works on *Synechocystis*, only the total PQ/PQH2 pool was measured, without distinguishing its photoactive fraction (Schuurmans et al. 2014). In *Chlorella*, VJ’ increased 1.06- and 1.21-fold after dark-to-light transition under high CO2 and air, respectively (Fig. 3). Comparative data for algae is scarce. PQ/PQH2 was measured in *Chlamydomonas reinhardtii* and *Arabidopsis thaliana*, however, the size of the photoactive pool shifted between low light and high light treatments (Pralon et al. 2019; Virtanen and Tyystjärvi 2023) - complicating estimation of the reference states.

Furthermore, in our experiments using Multi-Color PAM with a 725 nm LED, pretreatment of fully dark-acclimated cultures with far-red light never reduced VJ below the value of 0.46 in *Chlorella* and 0.61 in *Synechocystis* (data not shown). This is in a sharp contrast to many previous studies, in which far-red light was routinely applied to achieve full oxidation of the PQ pool. Moreover, far-red illumination enables estimation of the photoactive pool size, revealing that approximately 20–30% of the photoactive PQ pool remains reduced in *Synechocystis* and *Arabidopsis thaliana* under such conditions (Kruk and Karpinski 2006; Khorobrykh et al. 2020). These observations imply that the fluorescence method provides lower effective resolution than HPLC measurements for PQ-redox assessment.

Despite difficulties in the direct comparison with previous quantitative data, the experimental setup used in this study clearly shows that the VJ’ parameter can reflect PQ-redox changes on a semi-quantitative basis during dark-to-light transition (Figs. 3-4), light-to-dark transition (Supplementary Figure S6) as well as after reduction of PQ-pool by GA or KCN under actinic light (Figs. 5-6). During GA and KCN treatments, VJ’ reached the value of 1. The questions whether under those treatments, the whole photoactive PQ pool was reduced, or whether size of the photoactive pool changed, remains to be resolved.

As shown above, critical parameters for the OJIP fluorescence measurements are strong SP and low culture density (Tomek et al. 2001; Kumar Panigrahi and Kumar Mishra 2019), as well as relatively low AL and absence of PSII inhibitors (Figs. 2, 7). In order to reflect PQ-redox shifts between control and treated samples, identical sample pre-treatment is further required. In addition to pre-illumination or the presence or absence of inhibitors, this includes, for instance, constant temperature. In addition, VJ does not reflect PQ-redox well in samples with distinct Y(NA) states, as those alterations influence the IP phase (Supplementary Figure S9).

### Lessons for understanding PQ pool homeostasis in algae and cyanobacteria

Our results show that *Synechocystis,* in comparison with *Chlorella*, keeps impressively tight control over PQ-redox across a wide range of conditions, in agreement with previous works (Lea-Smith et al. 2013; Schuurmans et al. 2014). In *Synechocystis,* the PQ pool can become reduced either by exposure to high light, or upon cultivation under red light, even of very low intensity (Zavřel et al. 2024; Espinoza-Corral et al. 2024). To cope with high light, *Synechocystis* adopted several acclimation mechanisms such as reduction of the light-harvesting antenna and abundance of PETC components (Jahn et al. 2018), shifts in the PSII/PSI ratio (Kopecná et al. 2012) or an increase of intracellular glycogen (Zavřel et al. 2019), lipid and/or carotenoids (Zavřel et al. 2021). These strategies contribute to adjusting light energy distribution between photosystems, balancing PQ-redox and ATP to NADPH ratio (Höper et al. 2024). At the onset of high light exposure, the most important for maintaining PQ-redox homeostasis is PBS decoupling and activation of OCP-NPQ. In addition, as we also confirm here, PQ-redox is further balanced by TOs (Lea-Smith et al. 2013; Nikkanen et al. 2021). Indeed, also high CO2 helps to prevent PQ-pool overeduction upon high light exposure (Khorobrykh et al. 2020).

*Chlorella* possesses PTOX in its chloroplasts. However, this TO appears less effective in regulating PQ-redox compared to *Synechocystis*. Under the conditions of high energy inflow, increased PETC activity is accompanied by an elevated proton translocation and building up of a significant pH gradient across the thylakoid membrane, resulting in acidification of the thylakoid lumen. In *Chlorella*, lowering the lumenal pH induces xanthophyll cycle and NPQ, which, together with State I → State II transition, downregulates the amount of energy exciting P680 and reducing QA. The pH level in chloroplasts is under homeostatic control, which is, however, not yet fully understood (Trinh and Masuda 2022). The pH homeostasis in chloroplasts also contributes to the long-term regulation of PQ-redox balance. However, as shown also here, upon sudden shifts in light intensity, PQ-redox can face rapid changes since algae (similar to plants) have only limited options of its control.

### Conclusions

In this study, we critically assess the use of fast Chl *a* fluorescence rise kinetics (OJIP transients) as an indicator of the plastoquinone pool redox state (PQ-redox) in liquid cultures of green algae and cyanobacteria. We demonstrate that, despite certain limitations, the method is sufficiently sensitive to detect relative PQ-redox changes in both dark- and light-acclimated *Chlorella* and *Synechocystis* cells, using VJ and VJ’ parameters. The method requires 1) saturation pulse (SP) strong enough to induce rapid QA- accumulation; 2) low culture density to prevent fluorescence scattering and reabsorption; 3) similar rates of QA- accumulation in the control and treated samples and 4) unimpeded electron flow between QA-, QB and PQ. The VJ parameter reflects the PQ-redox best in dark-acclimated cultures. Under low to moderate actinic light, the VJ’ parameter is still able to reflect PQ-redox changes, however, not as reliably as in the dark. This is because part of QA gets reduced to QA- even before a saturation pulse is applied, resulting in less QA being available to form equilibrium with PQ. This implies that the VJ or VJ’ range, reflecting PQ-redox, strongly depends on the culture pre-treatment. Nevertheless, here we demonstrate that even under 100 µmol photons m^−2^ s^−1^ of white light, the VJ’ parameter reliably reflected substantial PQ-pool reduction induced by GA and KCN treatments, as well as PQ-pool oxidation induced by MV. Despite its semi-quantitative manner, the fluorescence method provides an easy-to-use alternative to traditional HPLC-based approaches for detecting PQ-redox shifts. The method, being fast and non-invasive, has a great potential to become widely used in algae and cyanobacteria research, in addition to plant research where it is already well established.

## Supporting information

Supplemental File with all supplementary figures and tables

## List of abbreviations

AL: actinic light
CBB: Calvin-Benson-Bassham cycle
COX: cytochrome *c* oxidase
Cyt *b*6/*f*: Cytochrome *b*6/*f* complex
Cyd: cytochrome *bd* quinol oxidase
DCMU: 3 (3,40-dichlorophenyl)-1,1-dimethylurea
FO, FO’: initial fluorescence yield at the beginning of the OJIP curve in fully dark-acclimated samples (≥ 20 min in darkness) and in light-acclimated or partially dark-acclimated samples, respectively
FI, FJ and FP: fluorescence yield at the J, I and P point of the OJIP curve in fully dark-acclimated cells (≥ 20 min in darkness), respectively
FI’, FJ’ and FP’: fluorescence yield at the J, I and P point of the OJIP curve in light- or partially dark-aclimated cells, respectively
FM, FM’: maximal fluorescence yield in fully dark-acclimated cells (≥ 20 min in darkness) and in light-acclimated or partially dark-acclimated samples, respectively
FNR: ferredoxin-NADP^+^-oxidoreductase
GA: glycolaldehyde
HL: high light
J, I, P: J, I and P points of the OJIP curve, respectively
ML: measuring light
MV: methyl viologen
NDH-1: NAD(P)H dehydrogenase-like complex type 1
NPQ: non-photochemical quenching
OEC: oxygen evolving complex of PSII
OCP: orange carotenoid protein
PBS: phycobilisomes
PC: plastocyanin
PFD: photon flux density
PQ: plastoquinone
PQH2: plastoquinol
PQ-redox: redox state of the PQ/PQH2 pool
PETC: photosynthetic electron transport chain
PSI: Photosystem I
PSI-CEF: cyclic electron flow around PSI
PSII: Photosystem II
PTOX: plastid terminal oxidase
QA: quinone A
QB: quinone B
SDH: succinate dehydrogenase
SP: saturating pulse
TOs: terminal oxidases
VJ, VJ’: relative fluorescence yield at the J point of the OJIP curve measured in fully dark-acclimated and in partially dark- or light-acclimated samples, respectively
Y(NA): quantum yield of non-photochemical energy dissipation due to acceptor side limitation of PSI

## Ethics approval and consent to participate

Not applicable

## Consent for publication

Not applicable

## Availability of data and materials

The datasets generated during and/or analysed during the current study are available in the Figshare repository under DOI <u>10.6084/m9.figshare.28938632</u>. The python scripts created and used to evaluate the measured OJIP curves can be accessed at https://github.com/Computational-Biology-Aachen/OJIP-PQredox. The repository includes a Jupyter notebook (paper_figures.ipynb) that reproduces all figures in the manuscript. Additionally, we provide a freely accessible online tool for streamlined processing and preliminary analysis of OJIP fluorescence data from the three fluorometers evaluated in this study, available at https://tools-py.e-cyanobacterium.org/.

## Authors’ contributions

TZ, AChP and TP designed and planned the research, performed the experiments and evaluated the data, TZ wrote the manuscript with contributions from all authors. All authors reviewed, revised and approved the final manuscript for publication.

## Acknowledgements

Not applicable

