## Supplemental File with all supplementary figures and tables for "Estimating the redox state of the plastoquinone pool in algae and cyanobacteria *via* OJIP fluorescence: perspectives and limitations"

**Supplementary Table S1** Overview of the settings of the three fluorometers used and the culture densities applied throughout the experiments described in this study.

| Strain | ***Chlorella*** | | | ***Synechocystis*** | | |
|---|---|---|---|---|---|---|
| Fluorometer | **MC-PAM** | **FL-6000** | **AquaPen** | **MC-PAM** | **FL-6000** | **AquaPen** |
| *Experiments evaluating the effect of saturation pules intensity and culture density on the OJIP transient (Fig. 1)* | | | | | | |
| Saturation pulse color (nm) | 440 | 460 | 455 | 625 | 623 | 630 |
| Saturation pulse intensity (μE $m^{-2}$ $s^{-1}$) | 1110 - 3710 | 1125 - 7500 | 1125 - 3750 | 1020 - 3250 | 975 - 6500 | 975 - 2600 |
| Saturation pulse length (ms) | 630 | 1000 | 2000 | 630 | 1000 | 2000 |
| Measuring light color (nm) | 440 | | | 625 | | |
| Measuring light intensity setting | 10 | | | 10 | | |
| Actinic light intensity (μE $m^{-2}$ $s^{-1}$) | n.a. | n.a. | n.a. | n.a. | n.a. | n.a. |
| Actinic light color | n.a. | n.a. | n.a. | n.a. | n.a. | n.a. |
| Fluorescence detection range (nm) | 665 - 710 | 690 - 730 | 667 - 750 | 665 - 710 | 690 - 730 | 667 - 750 |
| Offset setting | | 0.085 | | | 0.08 | |
| Gain setting | 2 | 0.25 | | 2 | 0.35 | |
| Damping setting | 1 | | | 1 | | |
| Measuring frequency (Hz) | 100000 | | | 100000 | | |
| Measuring time constant (μs) | 140 | | | 140 | | |
| Culture density: Chl *a* (mg $L^{-1}$) | 0.4 - 22.9 | 0.1 - 24.6 | 0.6 - 21.2 | 0.3 - 13.4 | 0.3 - 17.1 | 0.2 - 20.8 |
| *Experiments evaluating the effect of DCMU, light, MV, GA, KCN and high light on the OJIP transient (Fig. 2-7)* | | | | | | |
| Saturation pulse intensity (μE $m^{-2}$ $s^{-1}$) | 2244 | | | 1904 | | |
| Saturation pulse color (nm) | 440 | | | 625 | | |
| Saturation pulse length (ms) | 630 | | | 630 | | |
| Measuring light color (nm) | 440 | | | 625 | | |
| Measuring light intensity setting | 10 | | | 10 | | |
| Actinic light intensity (μE $m^{-2}$ $s^{-1}$) | 100 | | | 100 | | |
| Actinic light color | White | | | White | | |
| Fluorescence detection range (nm) | 665 - 710 | | | 665 - 710 | | |
| Gain setting | 3 | | | 3 | | |
| Damping setting | 1 | | | 1 | | |
| Measuring frequency (Hz) | 100000 | | | 100000 | | |
| Measuring time constant (μs) | 140 | | | 140 | | |
| Culture density: $OD_{680}$ in MC-1000 (r.u.) | 0.4 | | | 0.4 | | |
| Culture density: Chl *a* (mg $L^{-1}$) | 0.5 | | | 0.7 | | |

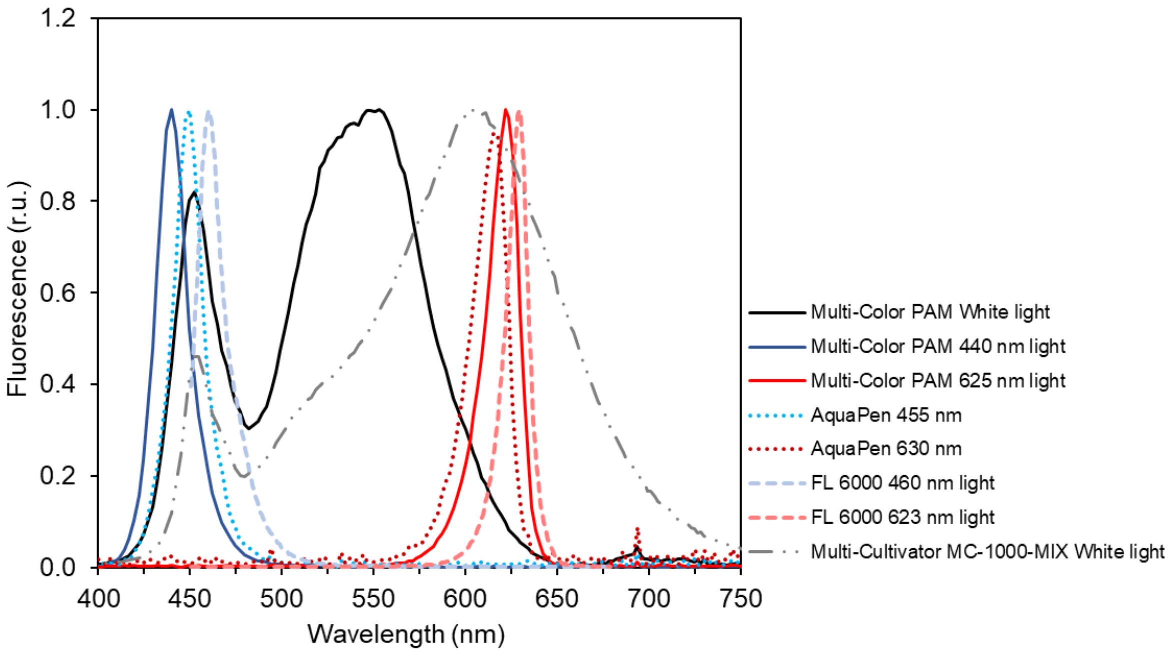


**Fig. S1** Emission spectra of white, blue and red LEDs in Multi-Color PAM (full black, blue and red lines, respectively), spectra of blue and red LEDs in AquaPen (dotted blue and red lines, respectively), spectra of blue and red LEDs in FL-6000 fluorometer (dashed blue and red lines, respectively), and spectra of white light LEDs in Multi-Cultivator MC-1000-MIX (dotdash black line). The spectra were recorded using the SpectraPen Mini (Photon System Instruments, Czechia) and normalized to their respective maxima prior to plotting.

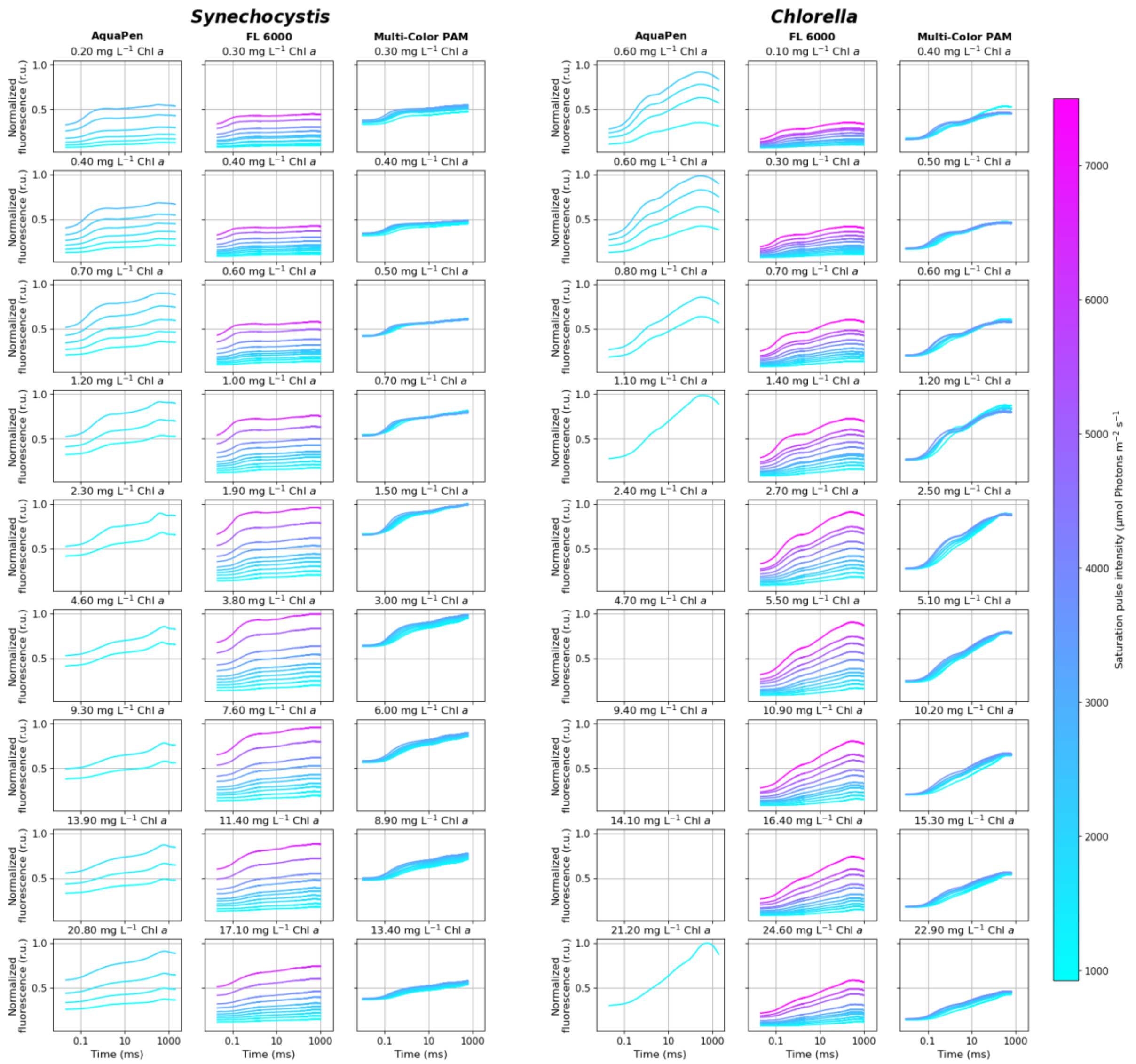


**Fig. S2** OJIP curves measured by FL 6000, Multi-Color PAM and AquaPen fluorometers across a matrix of SP intensity and culture density combinations. A common Y-axis scale is used, normalizing the raw OJIP curves to the maximal fluorescence values detected by each fluorometer. Missing values for AquaPen resulted from detector saturation; when the saturation occurred, the affected OJIP curves were excluded from analysis. Complete description of settings of the tested fluorometers, including SP intensities available, is provided in Supplementary Table S1.

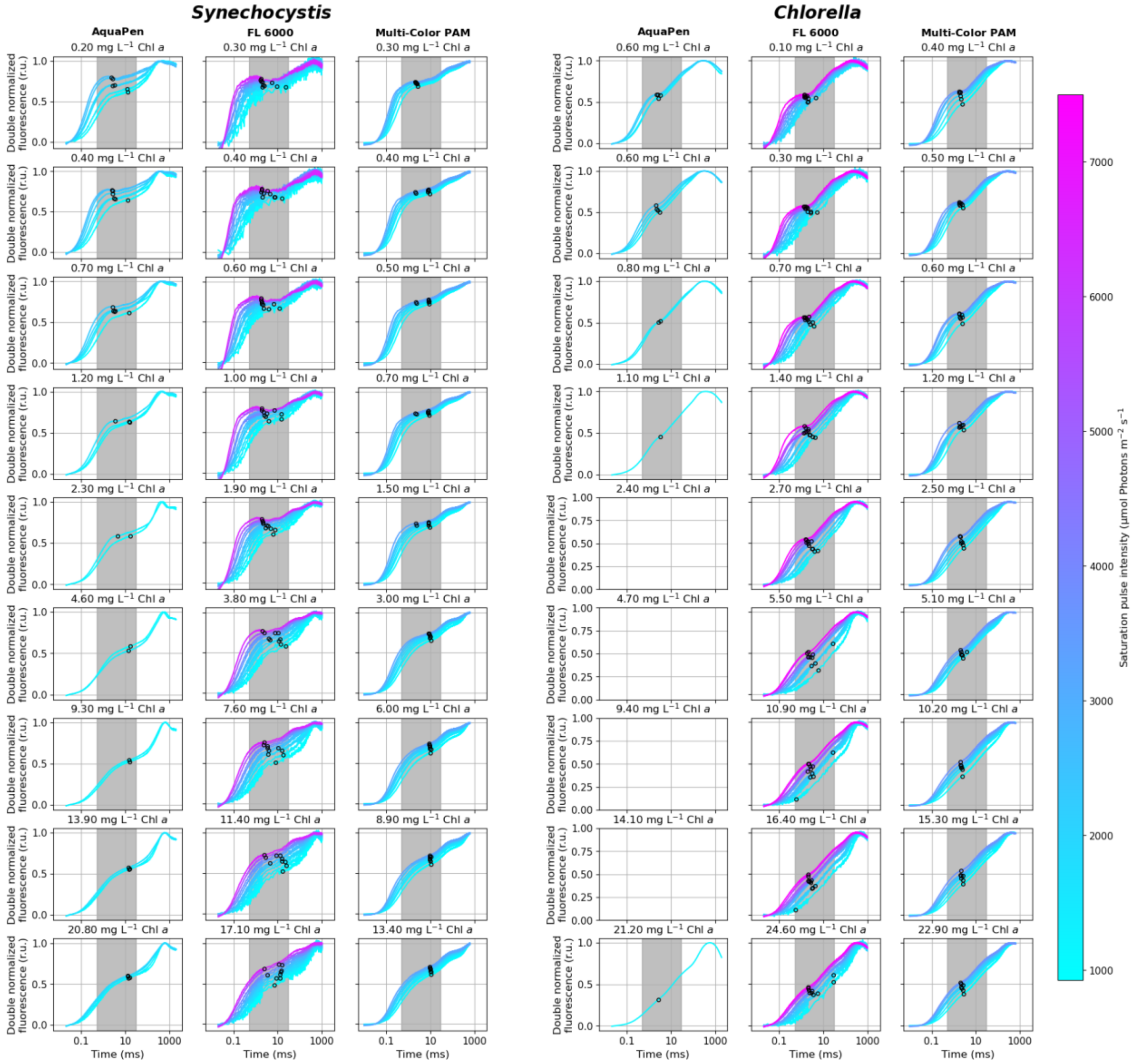


**Fig. S3** Double normalized OJIP curves from Fig. S1. The open circles represent timing of the first step of the fluorescence rise following $F_O$, determined by identifying inflection points within the 0.6 – 25 ms range, as marked by the grey rectangles. Missing values for AquaPen resulted from detector saturation; when the saturation occurred, the affected OJIP curves were excluded from analysis. Complete description of settings of the tested fluorometers, including SP intensities available, is provided in Supplementary Table S1.

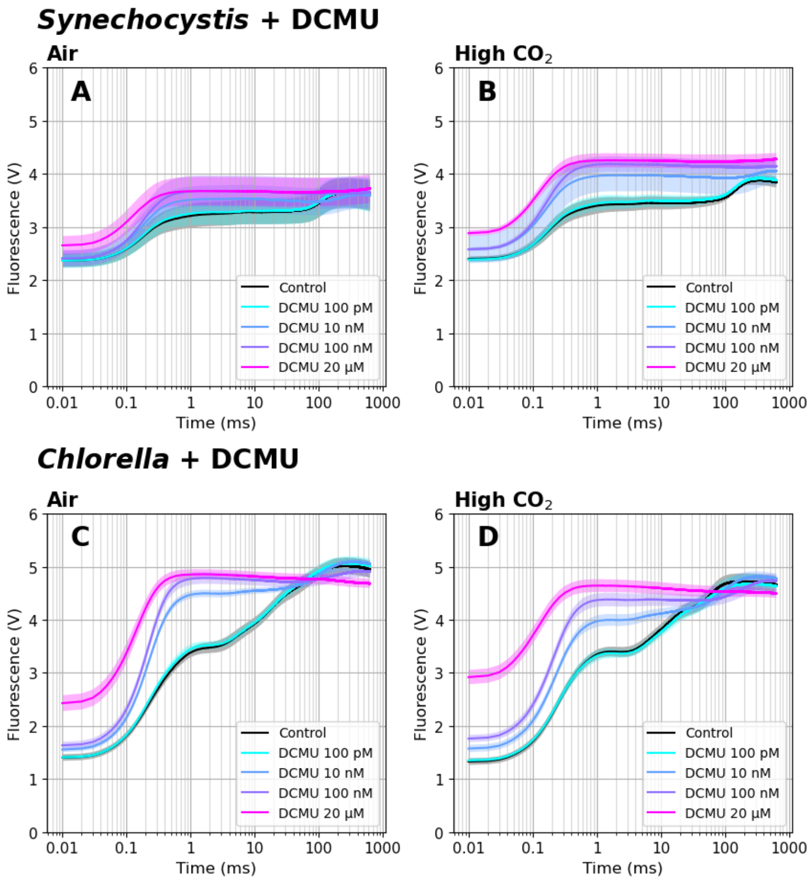


**Fig. S4** Raw fluorescence induction kinetics in the presence of 0 - 20 µM DCMU in *Synechocystis* (A-B) and *Chlorella* (C-D) cultures. Data represent averages ± SD, n = 4.

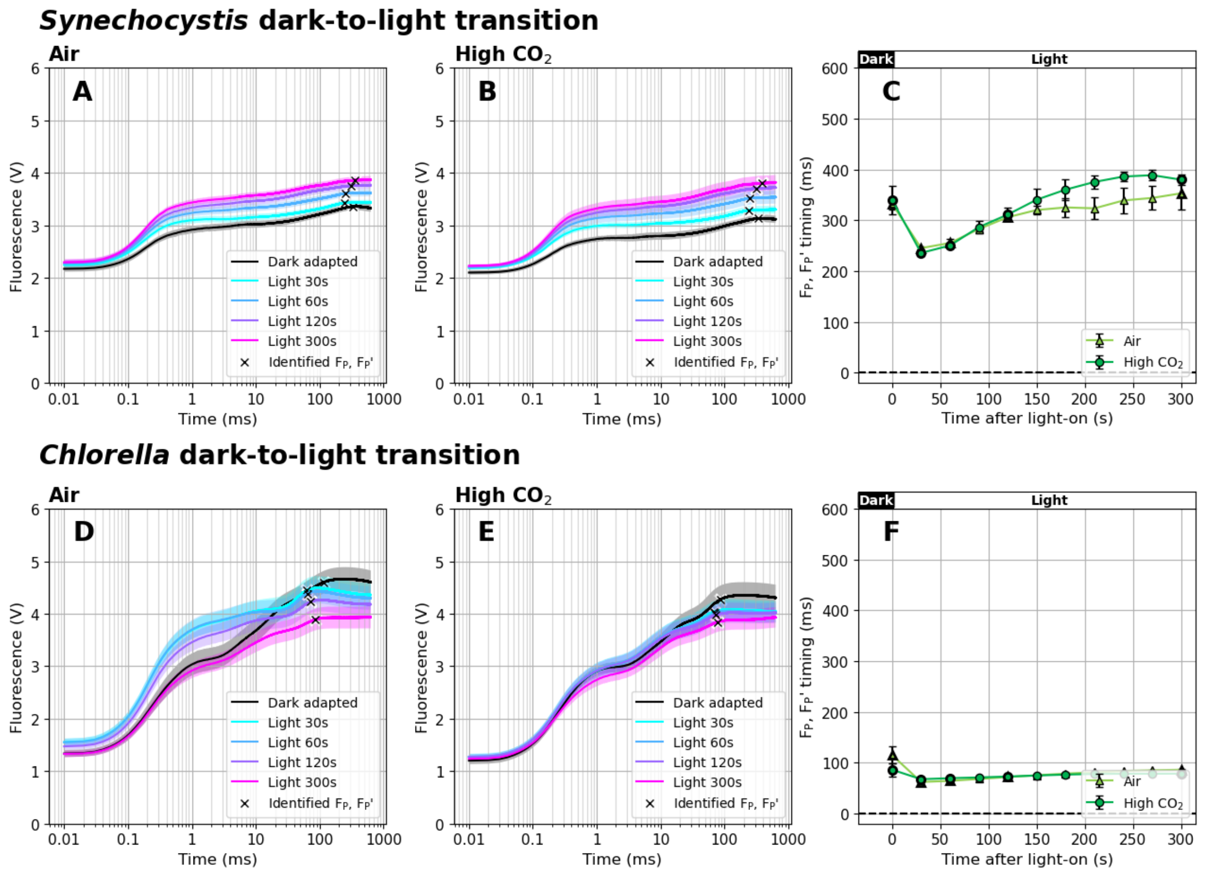


**Fig. S5** Raw fluorescence induction kinetics during dark-to-light transition in *Synechocystis* (A-B) and *Chlorella* (D-E) cultures. The timing of $F_P$ and $F_P$' (C, F), marked by crosses in panels A-B and D-E, was identified as a minimum of the second derivative of the fitted polynomial functions, approximating $F_P$ and $F_P$' as a fluorescence transient point closest to the expected appearance time (100 ms in *Chlorella*, 300 ms in *Synechocystis*). Data represent averages ± SD, n = 4.

**Light-to-dark transition**

For the light-to-dark transition experiments, *Synechocystis* and *Chlorella* cultures were cultivated under 10 µmol photons $m^{-2}$ $s^{-1}$ of white light in Erlenmeyer flasks that were sitting on a cultivation bench under ambient air atmosphere. For the measurement, 2 mL cultures aliquots were withdrawn from Erlenmeyer flasks and transferred to the cuvette of AquaPen (Photon System Instruments, Czechia). The fast fluorescence induction kinetics was measured immediately after the transfer (i.e. in light-acclimated state) and during 900 s of the dark acclimation.

In *Synechocystis*, $V_J$ was reduced to a lower level upon illumination and reached a steady state about three times faster compared to *Chlorella*, suggesting better control over PQ-redox (Supplementary Fig. S5). These results matched the expectations: it has been shown previously that FNR becomes inactivated (Schansker, Tóth, and Strasser 2006) and the PQ-pool becomes oxidized in the dark (Ivanov, Mubarakshina, and Khorobrykh 2007). The PQ-pool oxidation kinetics measured here (~120 s in *Synechocystis vs.* ~600 s in *Chlorella*, Supplementary Fig. S5) was remarkably faster than the kinetics of the FNR inactivation (~900 s, (Schansker, Tóth, and Strasser 2006)) but slower than previously reported kinetics of the PQ-pool oxidation (Ivanov, Mubarakshina, and Khorobrykh 2007). We note, however, that in the previous works either intact pea leaves, or isolated thylakoids from pea leaves were used. Variations of the determined rates can be thus expected.

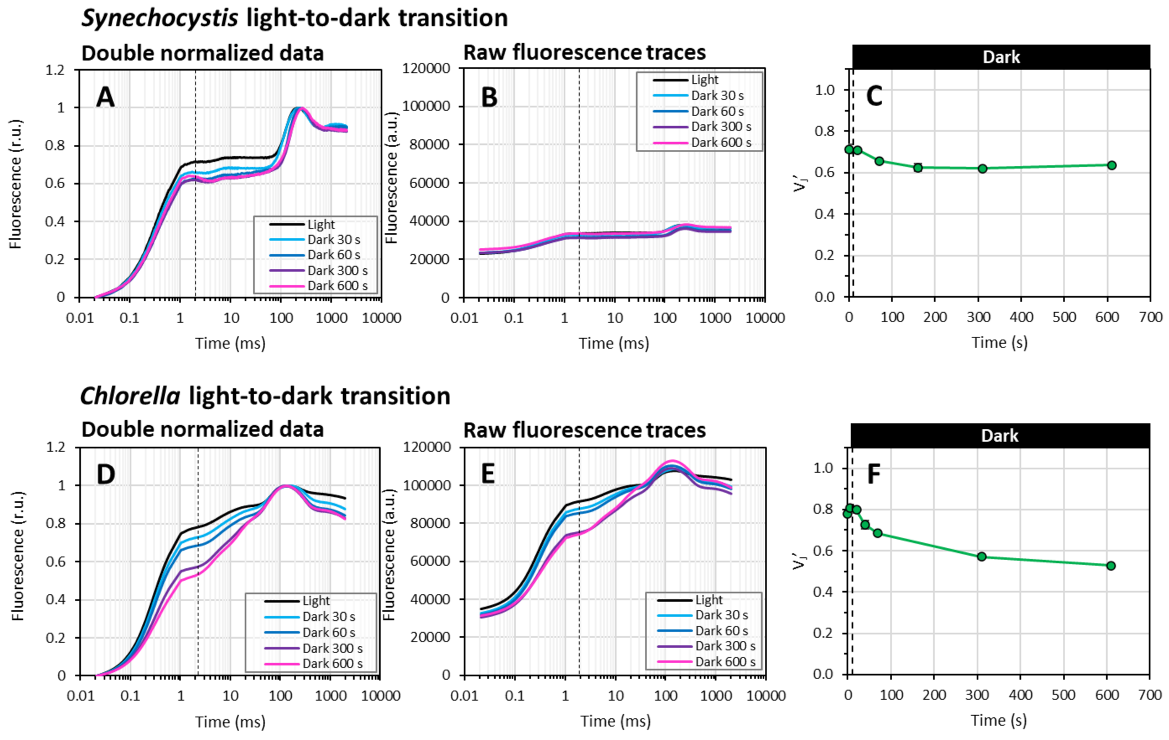


**Fig. S6** Fast fluorescence induction kinetics during light-to-dark transition. Cultures of *Synechocystis* (A-C) and *Chlorella* (D-F) were pre-cultivated in Erlenmeyer flasks under 10 µmol photons $m^{-2}$ $s^{-1}$ of white light on air. OJIP curves were recorded using an AquaPen (SP color: 625 nm (*Synechocystis*) / 455 nm (*Chlorella*) nm, SP PFD: 1 500 µmol photons $m^{-2}$ $s^{-1}$, SP duration: 2 000 ms) and the parameter $V_J$' (C, F) was calculated according to Eq. (1). Data represent averages ± SD, n = 3. Raw OJIP curves (A, D) and double normalized curves between $F_O$' and $F_M$' points (B, E) are shown without error bars for clarity. $F_J$' timing was identified as the first inflection point of the fluorescence signal around 2 ms. For simplicity, a single $F_J$' timing (marked by the dashed line in panels A-B and C-D) was used for the $V_J$' analysis within each treatment, corresponding with $F_J$' identified in the controlled culture.

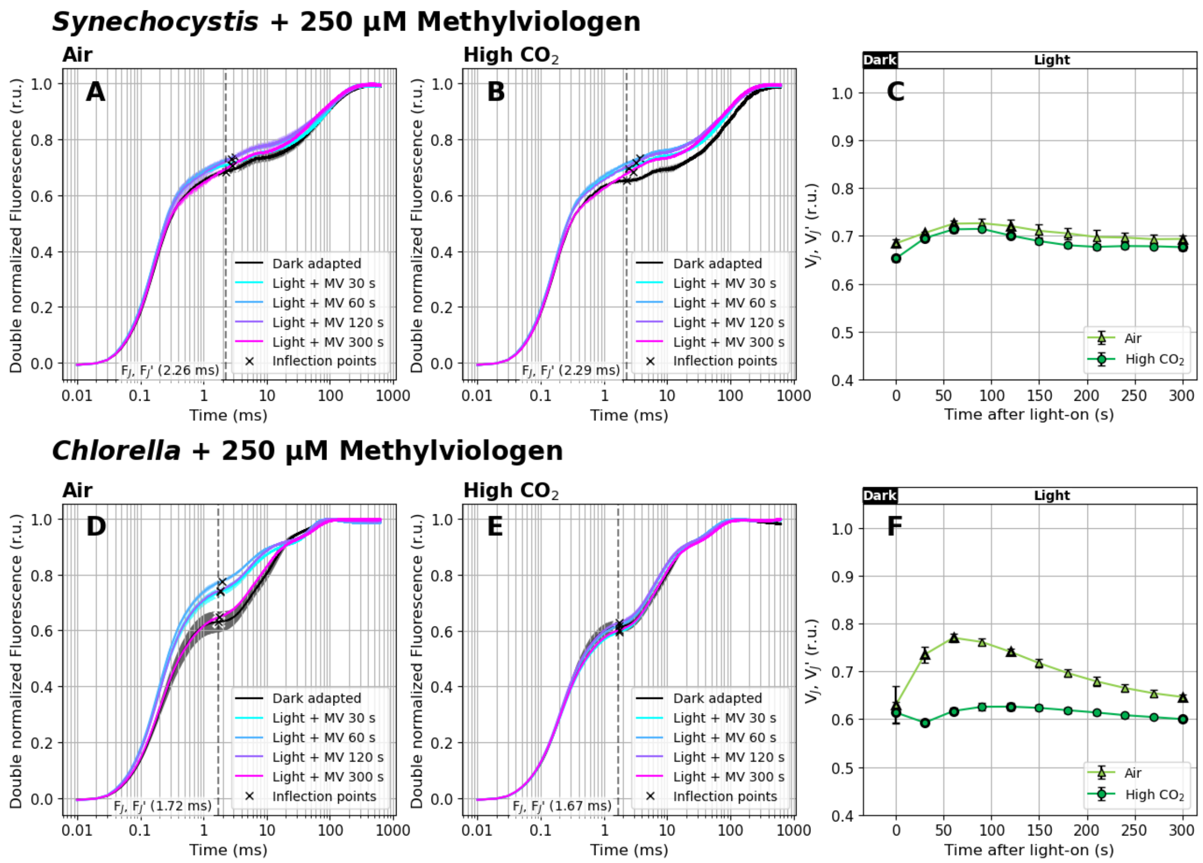


**Fig. S7** Double normalized fast fluorescence induction kinetics during dark-to-light transition in the presence of 0.25 mM methyl viologen (MV). Cultures of *Synechocystis* (A-C) and *Chlorella* (D-F) were pre-cultivated in a Multi-Cultivator under 25 µmol photons $m^{-2}$ $s^{-1}$ of white light under air (A, D) or 0.5 % $CO_2$ (B, E) and were dark-acclimated for 20 min prior to the fluorescence measurement in the presence of MV. After dark acclimation, OJIP curves were recorded using Multi-Color PAM and the parameter $V_J$ or $V_J$' (C, F) was calculated according to Eq. (1) and Eq. (2), respectively. OJIP curves in panels A-B and D-E were double normalized between $F_O$ and $F_M$ or between $F_O$' and $F_M$'. The raw OJIP curves are provided in Supplementary Fig. S6. $F_J$ and $F_J$' timing identified as the first inflection point of the fluorescence signal around 2 ms are marked by crosses in panels A-B and D-E. For simplicity, a single $F_J$ timing (marked by the dashed line) was used for the $V_J$' analysis within each treatment, corresponding with $F_J$ identified in the controlled culture. Data represent averages ± SD, n = 4.

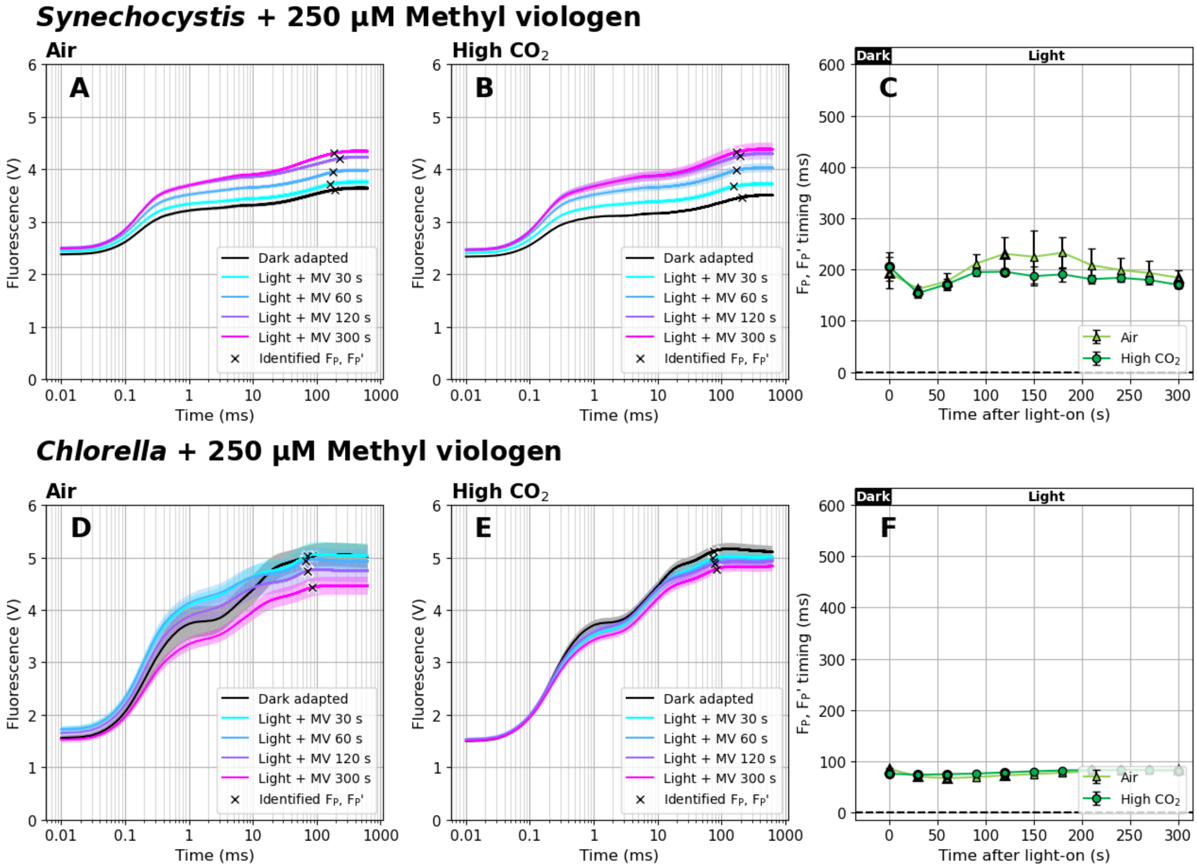


**Fig. S8** Raw fluorescence induction kinetics during dark-to-light transition in *Synechocystis* (A-B) and *Chlorella* (D-E) cultures in the presence of 0.25 mM methyl viologen (MV). The timing of $F_P$ and $F_P$' (C, F), marked by crosses in panels A-B and D-E, was identified as a minimum of the second derivative of the fitted polynomial functions, approximating $F_P$ and $F_P$' as a fluorescence transient point closest to the expected appearance time (100 ms in *Chlorella*, 200 ms in *Synechocystis*). Data represent averages ± SD, n = 4.

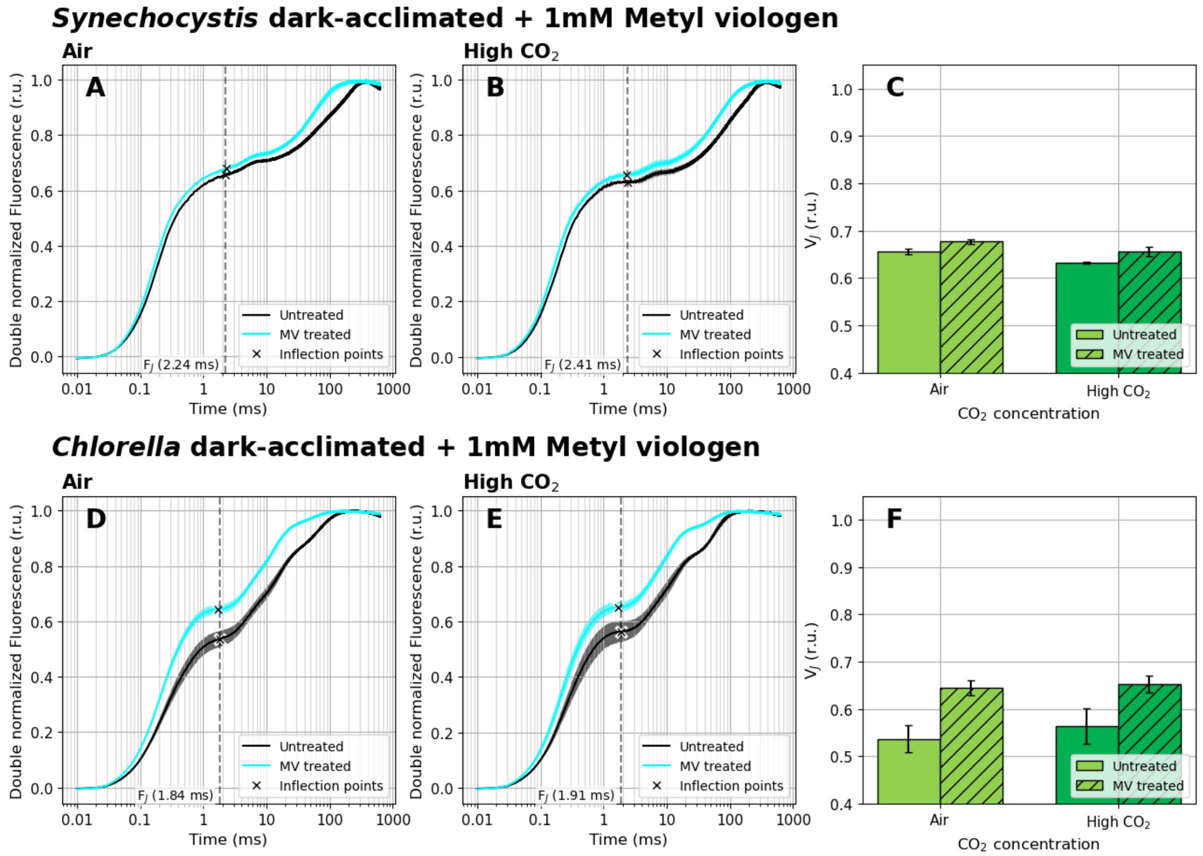


**Fig. S9** Double normalized fast fluorescence induction kinetics measured in dark-acclimated state in the absence or presence of 1 mM methyl viologen (MV). Cultures of *Synechocystis* (A-C) and *Chlorella* (D-F) were pre-cultivated in a Multi-Cultivator under 25 µmol photons m$^{-2}$ s$^{-1}$ of white light under air (A, D) or 0.5 % $CO_2$ (B, E) and were dark-acclimated for 20 min prior to the fluorescence measurement in the absence and presence of MV. After dark acclimation, OJIP curves were recorded using Multi-Color PAM and the parameter $V_J$ (C, F) was calculated according to Eq. (1). OJIP curves in panels A-B and D-E were double normalized between $F_{in}$ and $F_{max}$. The raw OJIP curves are provided in Supplementary Fig. S10. $F_J$ timing identified as the first inflection point of the fluorescence signal around 2 ms are marked by crosses in panels A-B and D-E. For simplicity, a single $F_J$ timing (marked by the dashed line) was used for the $V_J$ analysis within each treatment, corresponding with $F_J$ identified in the controlled culture (without MV). Data represent averages ± SD, n = 4.

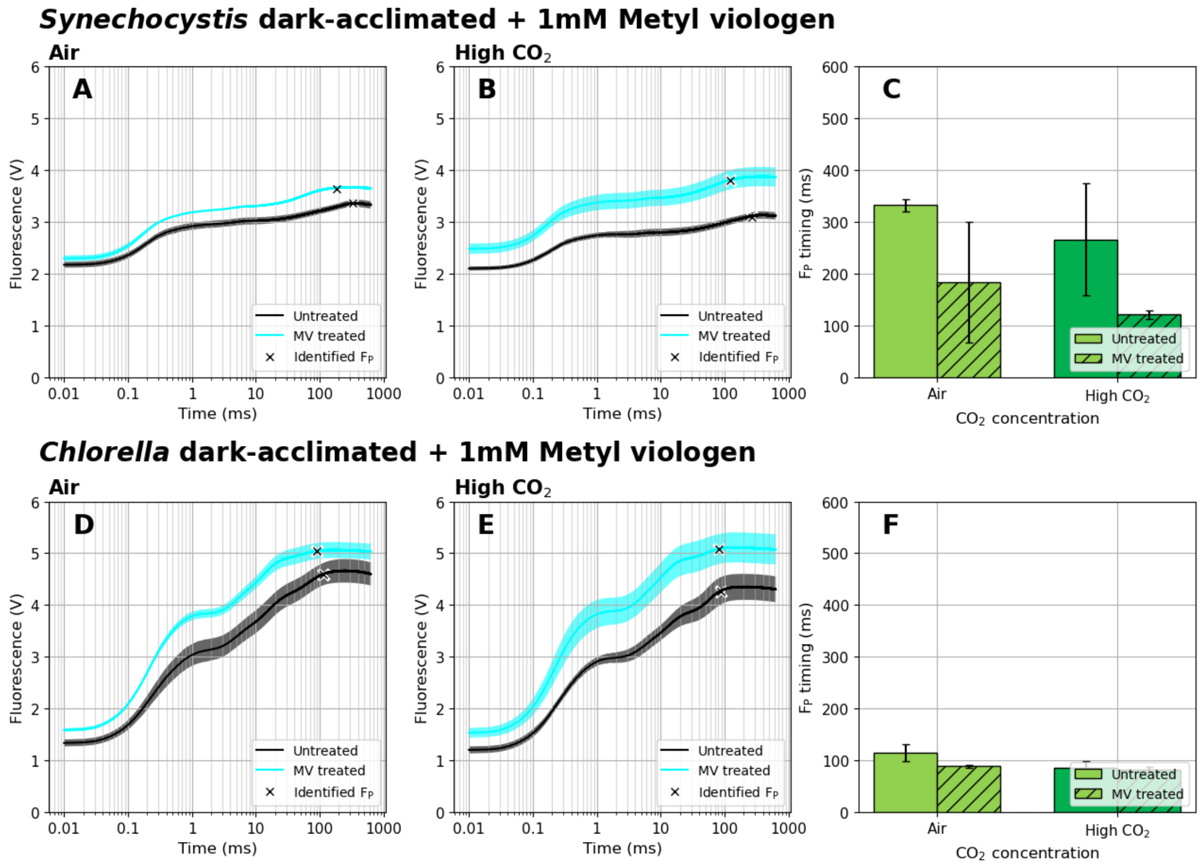


**Fig. S10** Raw fluorescence induction kinetics measured in dark-acclimated state in the absence or presence of 1 mM methyl viologen (MV) in *Synechocystis* (A-B) and *Chlorella* (C-D) cultures. The timing of $F_P$ (C, F), marked by crosses in panels A-B and D-E, was identified as a minimum of the second derivative of the fitted polynomial functions, approximating $F_P$ as a fluorescence transient point closest to the expected appearance time (100 ms in *Chlorella*, 200 ms in *Synechocystis*). Data represent averages ± SD, n = 4.

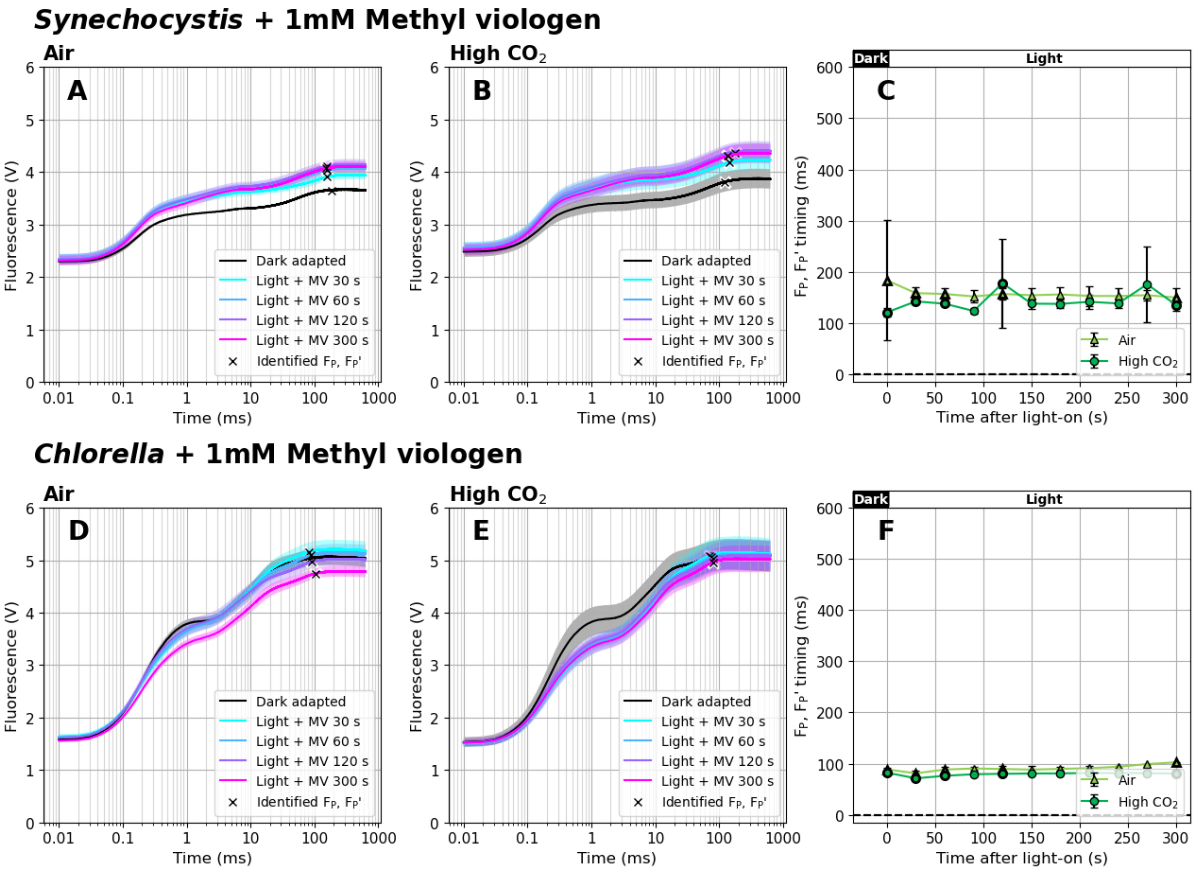


**Fig. S11** Raw fluorescence induction kinetics during dark-to-light transition in *Synechocystis* (A-B) and *Chlorella* (D-E) cultures in the presence of 1 mM methyl viologen (MV). The timing of $F_P$ and $F_P$' (C, F), marked by crosses in panels A-B and D-E, was identified as a minimum of the second derivative of the fitted polynomial functions, approximating $F_P$ and $F_P$' as a fluorescence transient point closest to the expected appearance time (100 ms in *Chlorella*, 200 ms in *Synechocystis*). Data represent averages ± SD, n = 4.

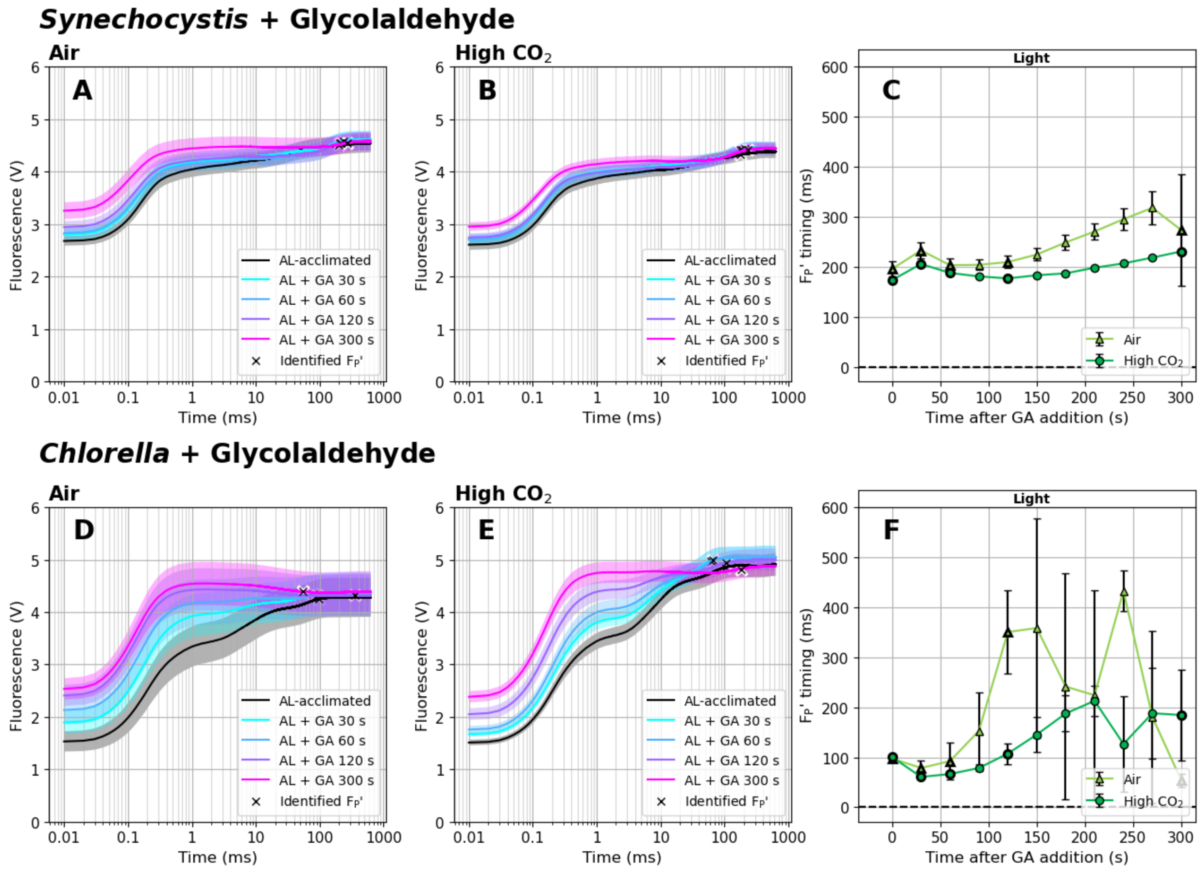


**Fig. S12** Raw fluorescence induction kinetics during dark-to-light transition in *Synechocystis* (A-B) and *Chlorella* (D-E) cultures in the presence of 25 mM glycolaldehyde (GA). The timing of $F_P$' (C, F), marked by crosses in panels A-B and D-E, was identified as a minimum of the second derivative of the fitted polynomial functions, approximating $F_P$' as a fluorescence transient point closest to the expected appearance time (100 ms in *Chlorella*, 200 ms in *Synechocystis*). Data represent averages ± SD, n = 4.

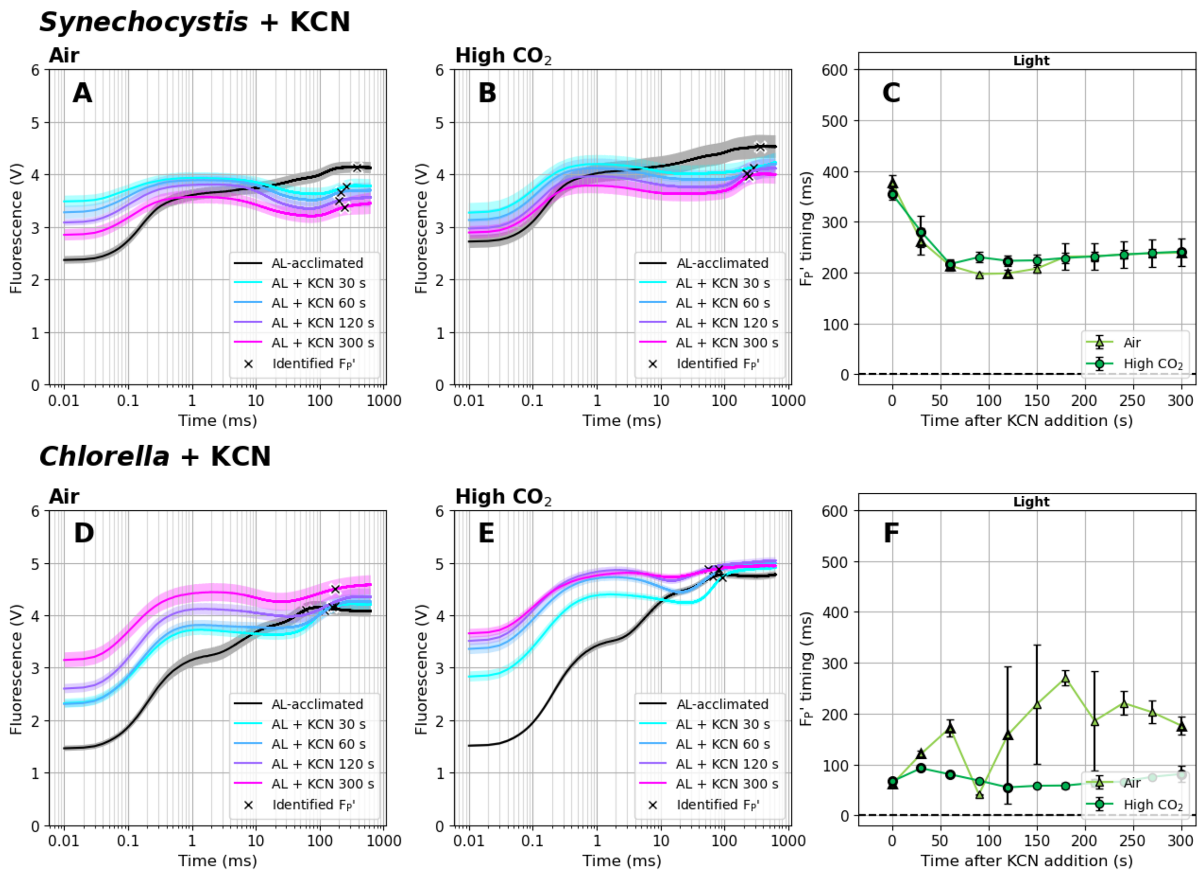


**Fig. S13** Raw fluorescence induction kinetics during dark-to-light transition in *Synechocystis* (A-B) and *Chlorella* (D-E) cultures in the presence of 1 mM potassium cyanide (KCN). The timing of $F_P$' (C, F), marked by crosses in panels A-B and D-E, was identified as a minimum of the second derivative of the fitted polynomial functions, approximating $F_P$' as a fluorescence transient point closest to the expected appearance time (100 ms in *Chlorella*, 300 ms in *Synechocystis*). Data represent averages ± SD, n = 4.

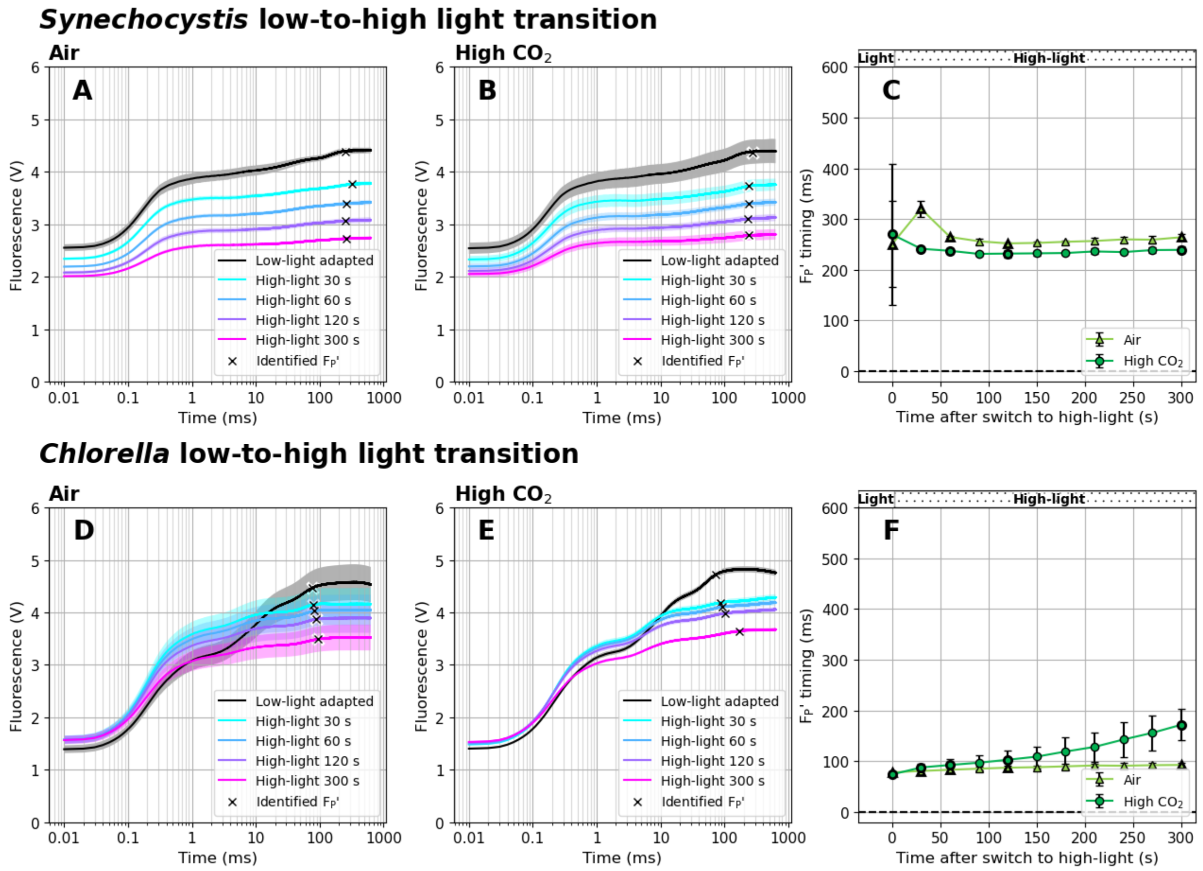


**Fig. S14** Raw fluorescence induction kinetics during high light treatment in *Synechocystis* (A-B) and *Chlorella* (D-E) cultures. The timing of $F_P$' (C, F), marked by crosses in panels A-B and D-E, was identified as a minimum of the second derivative of the fitted polynomial functions, approximating $F_P$' as a fluorescence transient point closest to the expected appearance time (100 ms in *Chlorella*, 300 ms in *Synechocystis*). Data represent averages ± SD, n = 4.

### High light treatment - shift in phycobilisome fractions

During an independent high light treatment experiment, cultures of *Synechocystis* were cultivated first in Multi-Cultivator and then in Erlenmeyer flasks in BG-11 medium under 60 µmol photons $m^{-2}$ $s^{-1}$ of white cultivation light at 23°C on air. High light (HL) treatment was applied by placing the flasks under 1 500 µmol photons $m^{-2}$ $s^{-1}$ of white light (supplemented by SANSI LED 15W light bulb). Before and during HL treatment, 5 mL culture aliquots were filtered under identical HL conditions (GF/B filters, Whatman, Maidstone, UK), flash-frozen in liquid nitrogen and stored at −80°C. Fluorescence excitation–emission maps were recorded at 77 K using a Jasco FP-8550 spectrofluorometer (Jasco, Tokyo, Japan). Right before the measurement, a slice from each filter (~1 cm × 0.3 cm) was cut to fit the metal holder of the transparent finger of the Dewar flask. 3D spectra maps were recorded over the excitation range 350–650 nm (step: 5 nm) and emission range 620–800 nm (step: 0.5 nm, bandwidth: 5 nm, scan speed: 1 000 nm $min^{-1}$, sensitivity: low). To distinguish fluorescence originating in phycobilisomes functionally attached to PSII (PBS-PSII) or PSI (PBS-PSI) and of phycobilisomes decoupled from both photosystems (PBS-free) the following equations were used (Zavřel et al. 2024):

$$PBS - free\ (normalized) = \frac{Ex_{360}\ Em_{661}}{PBS{-}total} \quad (S1)$$

$$PBS - PSII\ (normalized) = \frac{0.937 * Ex_{360}\ Em_{685} - 0.695 * Ex_{440}\ Em_{685}}{PBS{-}total} \quad (S2)$$

$$PBS - PSI\ (normalized) = \frac{0.937 * Ex_{360}\ Em_{726} - 0.695 * Ex_{440}\ Em_{726}}{PBS{-}total} \quad (S3)$$

$$PBS - total = PBS - free + PBS - PSII + PBS - PSI \quad (S4)$$

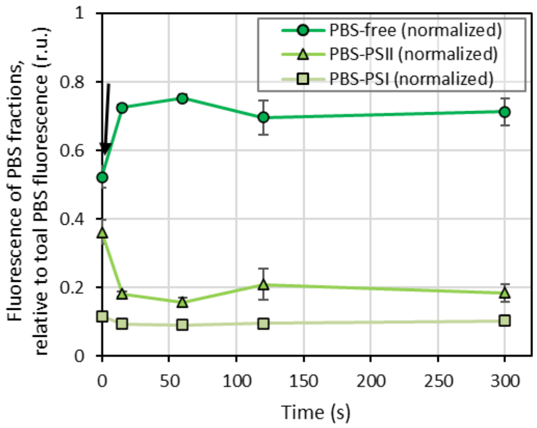


**Fig. S15** Shifts in fraction of phycobilisomes functionally attached to PSII (PBS-PSII), to PSI (PBS-PSI) and of phycobilisomes decoupled from both photosystems (PBS-free) during the high light (HL) treatment as described above. The timing of HL induction is marked by the black arrow. The parameters, derived from 77K spectra measurement, were calculated according to Eq. (S1-S4); the values represent averages ± SD, n = 3.
